# mTOR signaling governs the formation of epithelial apical projection via S6K1-RhoA and aPKC-Lgl2 axes

**DOI:** 10.1101/2024.12.08.627438

**Authors:** Sumit Sen, Prithwish Ghosh, Sudipta Mukherjee, Shreedha Prabhu, Ginni Khurana, Clyde Savio Pinto, Kirti Gupta, Ravindra Venkatramani, Mahendra Sonawane

**Affiliations:** Department of Biological Sciences, Tata Institute of Fundamental Research, Mumbai, India; Department of Chemical Sciences, Tata Institute of Fundamental Research, Mumbai, India

**Keywords:** mTOR signaling, microridges, actin cytoskeleton, RhoA signaling, aPKC signaling, *has/apkc*, aPKC phospho-regulation, cell polarity, apical projections, molecular dynamics simulation, zebrafish, epidermis

## Abstract

In metazoans, epithelia perform functions of absorption, diffusion, and secretion. The actin-based apical projections on the epithelial cells, contribute to these functions and are formed via cell-autonomous mechanisms that control cell polarity, intracellular transport, and the cytoskeleton. However, the cues that function upstream of these cell-autonomous regulators remain poorly known. Using microridges on zebrafish epithelial cells as a paradigm, we show that mTOR, a metabolic sensor, regulates the formation of apical projections. Mechanistically, mTORC1 controls the RhoA-ROCK activity via S6K1 to prevent the overactivation of Non-muscle Myosin II (NMII) to restrict microridge elongation. Furthermore, genetic, biochemical, and molecular dynamics simulation analyses reveal that mTORC2 regulates microridge pattern by modulating the activity of aPKC via its differential phosphorylation at two conserved sites. We propose that mTOR integrates the developmental and/or metabolic status of epithelial cells with cell autonomously acting RhoA and aPKC to regulate tissue wide formation of apical projections.

## Introduction

In metazoans, epithelial tissues act as semipermeable barriers between the environment and the internal milieu and perform additional absorptive or secretory functions essential for survival. To efficiently carry out these functions, the apical domain of epithelial cells is folded into membrane projections such as microvilli. Furthermore, an epithelial surface is coated with a mucus layer providing a physio-chemical barrier against allergens, pathogens, and shear stress (Sheng & Hasnain, 2022). Microridges are apical projections on squamous epithelial cells involved in mucus retention and abrasion resistance (Pinto et al., 2019; Sperry & Wassersug, 1976). They are evolutionarily conserved in the vertebrate lineage (Depasquale, 2018). Unlike other vertically growing apical projections such as microvilli or cilia, microridges grow laterally long with a relatively shorter height of about 300-500nm (Pinto et al., 2019). Due to their lateral growth, these protrusions form labyrinthine patterns that are highly variable between species (Fishelson, 1984; Sperry & Wassersug, 1976). How microridges grow laterally long and form these diverse patterns remain poorly understood.

Microridges are composed of branched F-actin and are further supported by cytokeratin filaments (Inaba et al., 2020; Pinto et al., 2019). Apical constriction of epithelial cells contributes to microridge elongation in a non-muscle myosin II (NMII)-dependent manner (van Loon et al., 2020, 2021). Molecular players such as Rho GTPases, Arp2/3, Formin, etc. were also shown to also be important for microridge formation (Lam et al., 2015; Pinto et al., 2019). Further, aPKC, a regulator of apicobasal polarity, has been shown to restrict the microridge length by controlling Lgl2 levels at the apical domain, where it functions as a pro-elongation factor by interacting with non-muscle myosin II (NMII) (Raman et al., 2016). However, it is unclear how the formation of microridges is coordinated across the entire epithelium during development.

The mechanistic Target Of Rapamycin (mTOR) is an atypical serine/threonine kinase of the Phosphatidylinositol 3-kinase-related kinases (PIKK) family of proteins. It forms two different complexes – mTOR complex 1 with Raptor and mLST8, and mTOR complex 2 with Rictor, mLST8, and mSin1 as interacting sub-units [Reviewed in (Panwar et al., 2023)]. mTORC1 primarily regulates cell growth and proliferation downstream to growth factors and nutrients (Ben-Sahra & Manning, 2017). In addition to its prominent role as a regulator of cellular metabolism, mTORC1 has been shown to influence actin cytoskeleton via regulating the expression of Rho GTPase proteins (Liu, Luo, et al., 2010). On the other hand, mTORC2 was originally identified as a rapamycin-insensitive complex with a distinct role in F-actin organization via regulators like Protein Kinase C α, a direct substrate of mTORC2 (Jacinto et al., 2004; Sarbassov et al., 2004). mTORC2 is activated by growth factors, membrane tension, and by its association with ribosomes (Battaglioni et al., 2022; Panwar et al., 2023). The mTOR pathway is conserved in eukaryotes and, amongst other functions, plays an important role in epithelial development. In the developing murine epidermis, mTORC1 promotes cell proliferation and differentiation whereas mTORC2 regulates the asymmetric division of cells (Ding et al., 2016). In zebrafish, mTOR signaling is essential for intestinal morphogenesis (Makky et al., 2007). Whether and how mTOR regulates the formation of apical membrane projections in the epithelial cells, remains unknown.

Here, we use the developing zebrafish epidermis as a model to report that mTOR signaling plays a crucial role in regulating microridge formation by preventing their premature elongation. We show that mTORC1 and its substrate S6K1 dampen the activity of RhoA, which is essential for microridge elongation via NMII. We further use a genetic approach and molecular dynamics simulation to elucidate differential regulation of aPKC structure and activity by mTORC2 via phosphorylations at two specific Threonine (Thr) residues. We propose that mTOR signaling integrates either developmental, growth factor, and/or metabolic signaling with cell-intrinsic mechanisms to coordinate microridge formation across the epithelial tissue during development.

## Results

### Inhibition of mTOR function leads to precocious elongation of microridges

Given the importance of mTOR signaling in the regulation of metabolism, cell proliferation, cell growth, and cytoskeleton, (Panwar et al., 2023; Szwed et al., 2021) we asked whether this pathway plays any role in the formation of microridges. To test this, we treated zebrafish embryos with Rapamycin, a known inhibitor of mTOR, from 24 to 48 hours post fertilization (hpf). Consistent with earlier results, we did not observe any obvious morphological defects in the developing bi-layered epidermis (Gupta et al., 2022). We verified the efficacy of the inhibitor by assessing the phosphorylation of Ribosomal Protein S6 (RPS6) at Ser240/244 (fig. 1A), and of Akt at Ser473 (fig. 1C), downstream targets of mTORC1 and mTORC2 signaling, respectively. Rapamycin treatment significantly reduced the levels of both phosphorylated RPS6 and Akt in the periderm cells (fig. 1A-D; supp fig. 1A). We quantified the length of microridges at 48hpf using a custom-built algorithm (see methods for details) and plotted both distributions of the microridge lengths from all the cells that were quantified and the average length of microridges for individual cells. The quantification revealed that Rapamycin treatment led to significant elongation of microridges compared to vehicle-treated embryos (fig. 1E-G). Given that microridges become developmentally long after 72 hpf (Raman et al 2016), mTOR inhibition seems to result in a precocious elongation of microridges.

**Figure 1.**
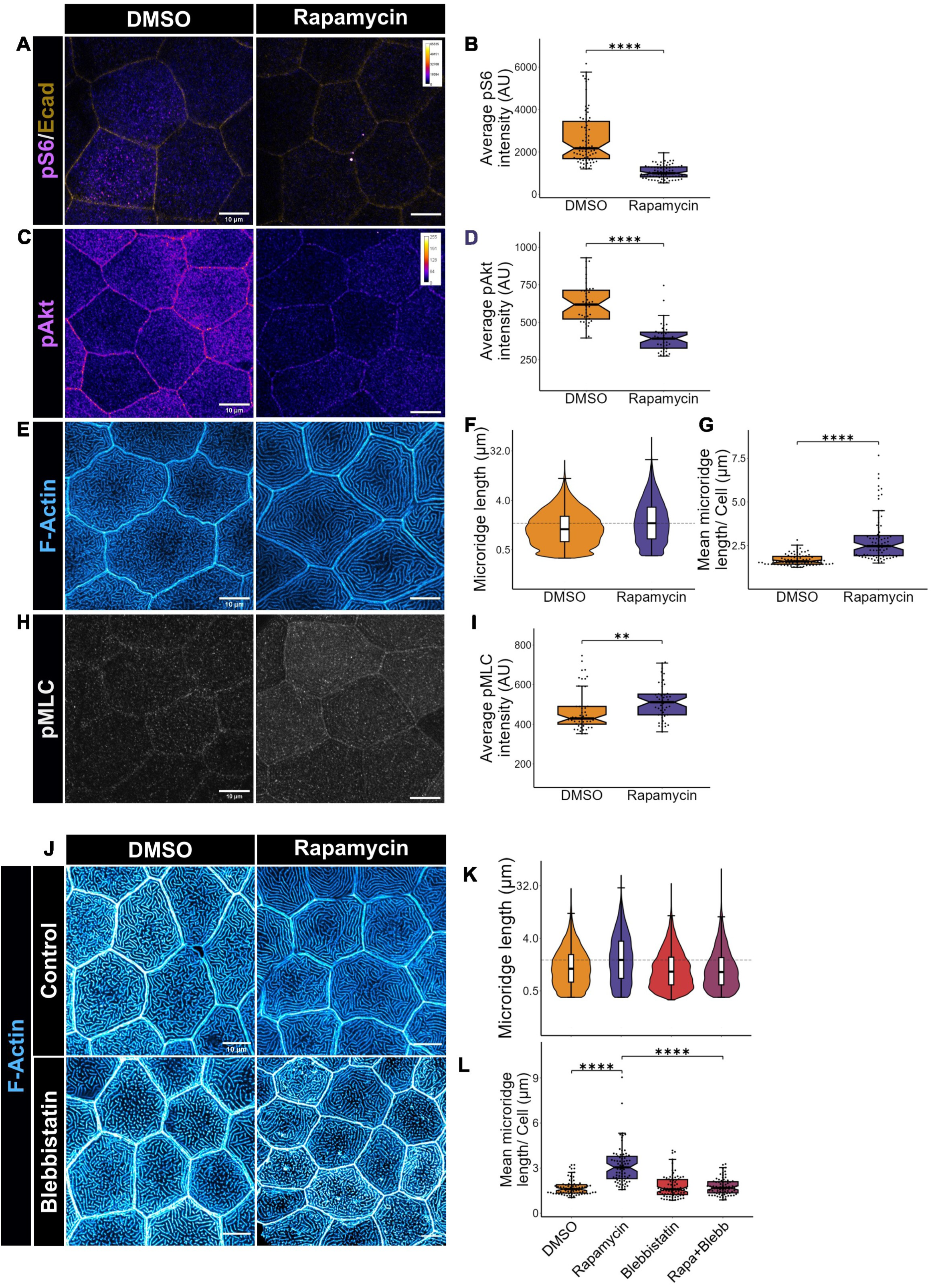
mTOR activity is essential to prevent NMII dependent precocious elongation of microridges. Confocal micrographs (A, C, E, and H) of peridermal cells in wildtype embryos at 48hpf treated with rapamycin and vehicle control (DMSO) starting at 24 hpf and immunostained for phospho RPS6 (Ser 240/244) (A), phospho Akt (Ser 473) (C), F-actin (E), and phospho MLC (Ser 19) (H). The confocal images (A, C) are represented in Fire LUT and the calibration bar ranges from 0-65535 indicating lowest to highest intensity. The graphs (B, D, and I) show the quantification of the average intensity of pS6 (B) and, pAkt (D) and pMLC (I). Violin plot represents microridge length distribution (F) and box-jitter plot of distribution of mean microridge lengths (G) in DMSO and rapamycin-treated embryos. (J) Rhodamine phalloidin staining of 48hpf embryos treated with rapamycin for 24hpf or blebbistatin (50μM, from 46.5-48 hpf) or both. (K) and (L) shows the quantification of these conditions as microridge length distribution and difference in mean microridge lengths, respectively. All the experiments were performed three times independently. For data represented in (B, F, G, K, and L) 75 cells from 15 embryos from three sets, for (D) 40 cells from 12-13 embryos, and (I) 50 cells from 10 embryos from two sets were analyzed. The third set for these two conditions imaged at 16 bit depth are presented in supp fig. 1. The asterisks in the plot represent significant differences between conditions. Mann-Whitney U test was used for (B, D, G, and I). For (L) Kruskal Wallis test with Dunn’s post hoc was used for multiple comparisons. Dotted lines in (F and K) represent highest median between the conditions. ** p<01; ****p<0.0001. The scale bar corresponds to 10μm.

Microridge elongation is driven by the activity of Non-Muscle Myosin II (NMII) (Raman et al., 2016; van Loon et al., 2021). To check whether the precocious elongation of microridges observed upon mTOR inhibition is correlated with increased NMII activity, we assessed the phosphorylation of myosin light chain (MLC) using a phospho-specific antibody made against pS19, an indicator of active NMII. In rapamycin-treated embryos, we observed a significant increase in pMLC levels (fig. 1H, I; supp fig. 1B) suggesting that myosin activity was enhanced upon mTOR inhibition. To further test whether NMII activity is responsible for microridge elongation upon inhibition of mTOR signaling, we treated embryos with Blebbistatin (an inhibitor of NMII) from 46.5 hpf to 48 hpf. The microridges were shorter in embryos with combined treatment of Rapamycin and Blebbistatin, as compared to those treated with just Rapamycin (fig. 1J-L). To further consolidate the importance of NMII activity in microridge elongation upon mTOR inhibition, we dampened the activity of Rho-associated protein Kinase (ROCK) using a pharmacological inhibitor Rockout (Yarrow et al., 2005). ROCK phosphorylates and activates the regulatory light chain of NMII (Riento & Ridley, 2003). Inhibition of ROCK activity led to shorter microridges in control as well as Rapamycin-treated embryos, thereby suggesting an essential requirement of ROCK activity for microridge elongation and pattern formation (fig. 2A-C).

**Figure 2.**
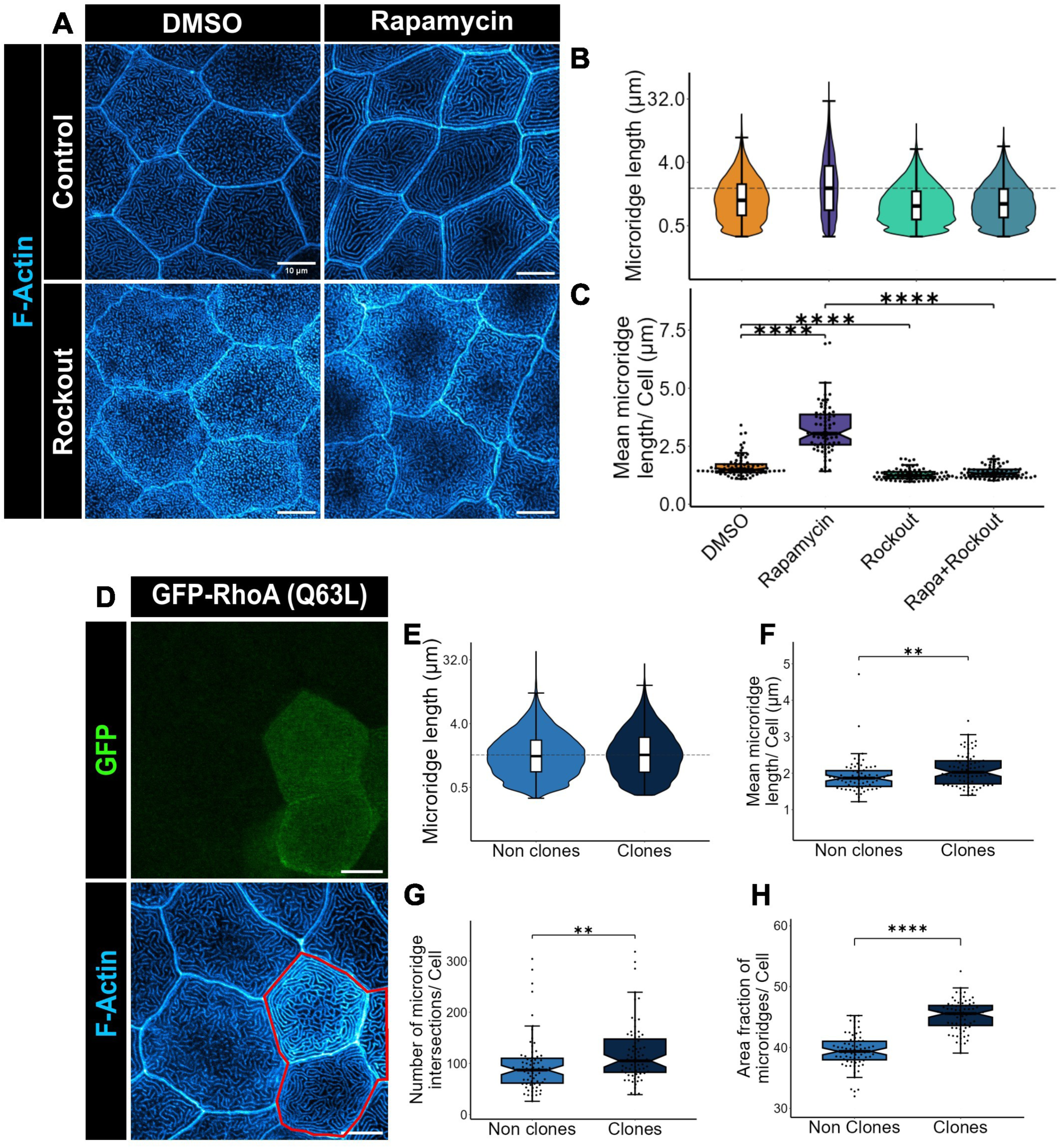
mTOR regulates microridge elongation via RhoA/ROCK pathway. Representative confocal scan of Rhodamine-phalloidin stained peridermal cells (A), distribution of microridge lengths (B) and box-jitter plot of mean microridge lengths of embryos treated with vehicle control (DMSO), rapamycin, Rockout (100μM, from 46.5-48 hpf), or both. (D) Representative image of peridermal cells from an embryo injected with GFP tagged constitutively active RhoA (Q63L) plasmid at one cell stage and stained at 48 hpf for GFP to identify CA-RhoA expressing cells and for F-actin using phalloidin. (E-H) shows quantification of different microridge parameters such as length distributions (E), mean microridge lengths (F), number of microridge intersections (G), and area covered by microridges in the apical domain (H) in the cells expressing CA-RhoA-GFP (clones) and their neighboring cells (nonclones). All the experiments were performed three times independently. For data presented in (B, C) 75 cells from 15 embryos and for (E-H) 71 cells from 26 embryos were analyzed. Asterisks in (C) show statistically significant difference using Kruskal Wallis test with Dunn’s post hoc test for multiple comparisons, while in (F, G, and H) the Mann Whitney U test was used. ** p<0.01, **** p<0.0001. The scale bar corresponds to 10μm.

Typically, ROCK is locked in an autoinhibited condition, which is later disinhibited by an active RhoA (Riento & Ridley, 2003). To confirm that RhoA activity regulates microridge elongation and pattern, we clonally expressed a constitutively active (CA) form of RhoA (GFP-RhoA-Q63L) in wild-type embryos (fig. 2D) by injecting plasmid as described before (Gupta & Sonawane, 2020). The cells expressing GFP-tagged CA-RhoA showed long and highly convoluted microridges with more interconnections and higher density than the neighboring non-clone cells (fig. 2D-H).

We conclude that mTOR signaling controls microridge elongation by suppressing the activity of the RhoA-ROCK-NMII axis in the squamous epithelial cells of the developing zebrafish epidermis.

### mTORC1 regulates microridges via the S6K1-RhoA axis

The two mTOR complexes, mTORC1 and mTORC2, have been shown to regulate RhoA by controlling its expression or activity, respectively (Liu, Das, et al., 2010; Liu, Luo, et al., 2010). To determine which of these two complexes regulates RhoA *in vivo* in zebrafish, we utilized the mechanism of action of rapamycin. Rapamycin was originally reported as a specific inhibitor of mTORC1. Eventually, it was shown that rapamycin also inhibits mTORC2 function upon longer incubation (Loewith et al., 2002; Sarbassov et al., 2006). Therefore, starting at 24 hpf, we assessed the effect of rapamycin on microridges at 2, 4, 6, and 12 hours post-treatment. We observed that within 2 hours of rapamycin treatment, mTORC1 activity diminished in the peridermal cells as assessed using pS6 staining (fig. 3A, B). In contrast, it required at least 6 hours for pAKT levels to decrease, which is a readout for mTORC2 activity (fig. 3C, D; supp fig. 1C). We also observed that microridges started to elongate within 4 hours of rapamycin treatment (fig. 3E-G), suggesting that the downregulation of mTORC1 is the primary cause of the early microridge elongation.

**Figure 3.**
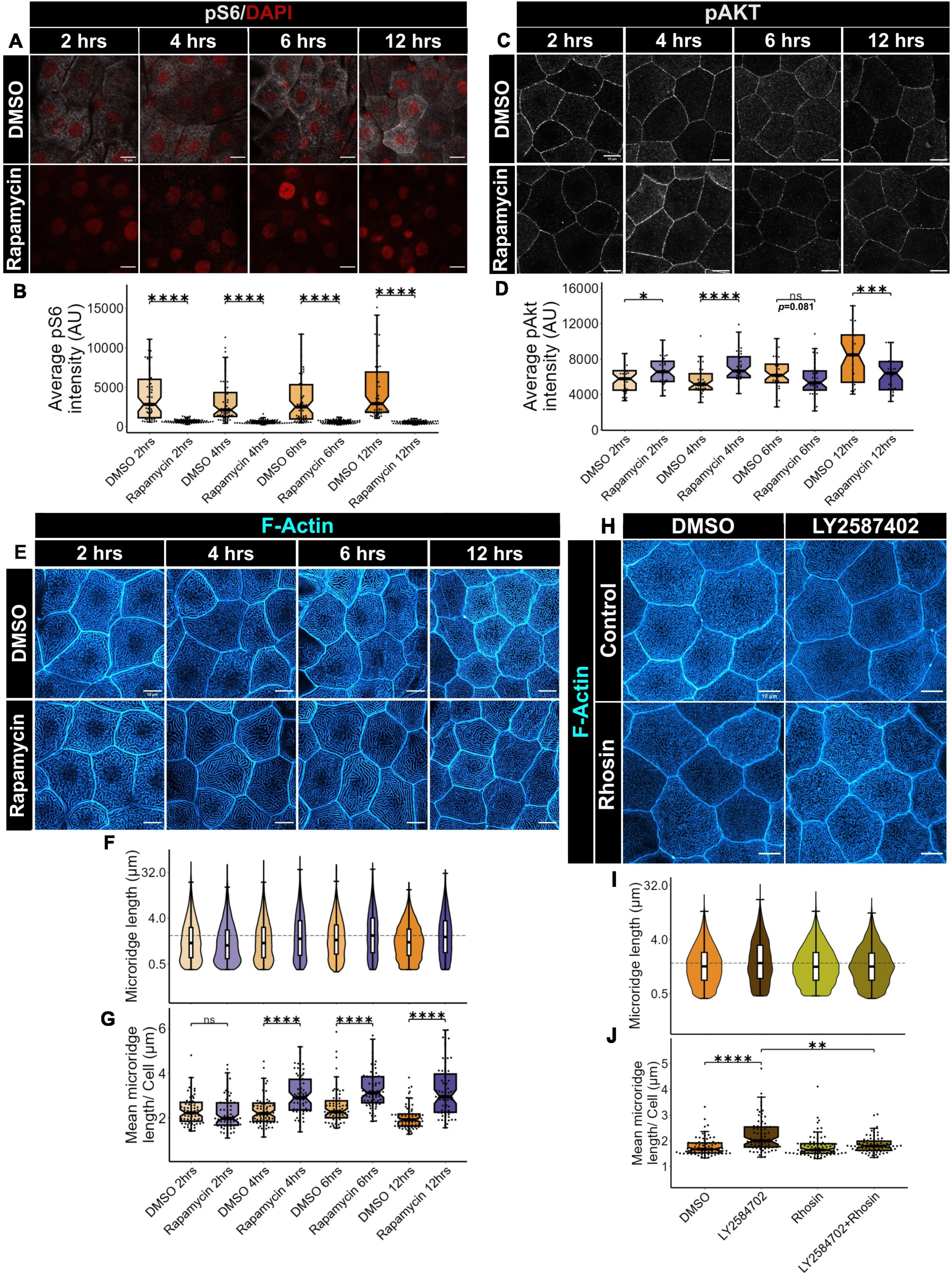
mTORC1/S6K1 pathway regulates RhoA activity in the zebrafish periderm. Representative images of peridermal cells of embryos treated with DMSO or rapamycin for 2, 4, 6, and 12 hours starting at 24hpf and immune-stained for pS6 (Ser 240/244) (A) and pAkt (Ser473) (C) and their corresponding intensity quantifications in (B) and (D), respectively. Confocal micrographs of microridges (E) in embryos treated with rapamycin or DMSO for the given time points starting from 24hpf and stained with rhodamine phalloidin and graphical representation of the distribution of microridge lengths (F) and mean microridge lengths (G) at these time points under given conditions. (H) Confocal images of rhodamine phalloidin stained of embryos treated with LY2587402 (S6K1 inhibitor) (10 μM), rhosin (40μM), or both at 48 hpf and graphical representations of the distribution of microridge lengths (I) and the mean microridge lengths (J) under these conditions. All the experiments were performed three times independently. For experiments (B, F, G, I, and J) 75 embryos from 15 embryos were analyzed. For (D) 39-46 embryos from 12-13 embryos were analyzed from two sets and the data for the third set, imaged at 12 bit depth, is presented in supp fig. 1. Asterisks in the plot represent significant differences in p-value analyzed by Mann Whitney U test in (B, D, and G) and Kruskal Wallis test followed by Dunn’s post hoc test for multiple comparisons in (J). *ns* (non significant) p>0.05, ** p<0.05, *** p<0.001, **** p<0.0001. The scale bar corresponds to 10μm.

To further assess whether RhoA function is required downstream of mTORC1, we used Rhosin - an inhibitor of RhoA activation (Shang et al., 2012) - in conjunction with the Rapamycin treatment. We observed that while the microridges elongate in the Rapamycin-treated embryos, Rhosin treatment along with Rapamycin for 4 hours, suppressed the elongation of microridges (supp fig. 1D-F), suggesting that RhoA activity is essential for microridge elongation upon mTORC1 inhibition. We further asked whether S6 Kinase 1 (S6K1) regulates RhoA activity. S6K1 is one of the downstream substrates of mTORC1, which regulates the actin cytoskeleton via directly binding to F-actin or regulating the activity of Rho GTPases, Rac1, and Cdc42 (Ip et al., 2011). To test the involvement of S6K1, we used an S6K1-specific ATP competitive inhibitor called LY2584702 (Tolcher et al., 2014). The embryos at 24 hpf, treated with LY2587402 for 24 hours, lowered the activity of S6K1 (supp fig. 2A, B) and displayed a long microridge phenotype similar to rapamycin treatment (supp fig. 2C-E). This microridge elongation is also associated with an increased NMII activity revealed by pMLC immunostaining (supp fig. 2F, G). Given this, we asked whether increased microridge elongation upon S6K1 inhibition is RhoA dependent. To test this, we treated embryos at 24 hpf with Rhosin along with the S6K1-specific inhibitor for 24 hours. Inhibition of RhoA activity under S6K1 downregulated condition rescued the microridge length (fig. 3H-J) suggesting that increased RhoA activation leads to microridge elongation downstream to S6K1 inhibition.

Our data suggest that mTORC1 restricts microridge elongation via its substrate S6K1 and RhoA signaling.

### mTORC2 regulates microridges by phosphorylating aPKC at turn motif

In the past, we have shown that the microridges elongate precociously upon the loss of aPKC (PKCλ/ɩ) function (Raman et al., 2016). This phenotype is similar to that observed upon mTOR inhibition. Therefore, we hypothesized that both aPKC and mTOR function in the same pathway to mediate microridge formation. To test this hypothesis, we first asked whether mTOR regulates the expression of the apkc (*prkci)* gene. A qPCR analysis on cDNA from embryos treated with rapamycin and a vehicle control revealed that in the rapamycin-treated embryos, *prkci* expression was significantly downregulated (supp fig. 3A). Surprisingly, the aPKC protein localization did not decrease in the peridermal cells upon 24 hours of rapamycin treatment (supp fig. 3B, C) suggesting that aPKC protein levels are maintained in these cells despite a reduction in *prkci* mRNA levels in the embryo. Given there was no change in aPKC levels upon rapamycin treatment, we asked whether aPKC is post-translationally modified by mTOR. aPKC belongs to the AGC kinase family proteins, and the turn motif of these proteins is phosphorylated by mTORC2 (Li & Gao, 2014; Tobias et al., 2016). In MEF cells, the turn motif of PKCζ is phosphorylated by mTORC2, which is required for downstream F-actin cytoskeleton regulation (Li & Gao, 2014). In zebrafish, this aPKCλ/ɩ residue is also conserved (fig. 4A). Since the aPKC λ/ɩ isoform expresses prevalently in the peridermal cells as previously reported (supp fig. 3D, E) (Cokus et al., 2019), we asked whether Thr (556) of turn motif is phosphorylated by mTORC2 in zebrafish. Using an aPKC λ/ɩ phospho-specific antibody, we found that the zf-aPKC λ/ɩ isoform is phosphorylated *in vivo* at the turn motif. Further, a prolonged rapamycin treatment of 24 hours, which downregulates mTORC2 (fig. 1F, G), decreased the level of phosphorylation at thr556 (fig. 4B, C).

**Figure 4.**
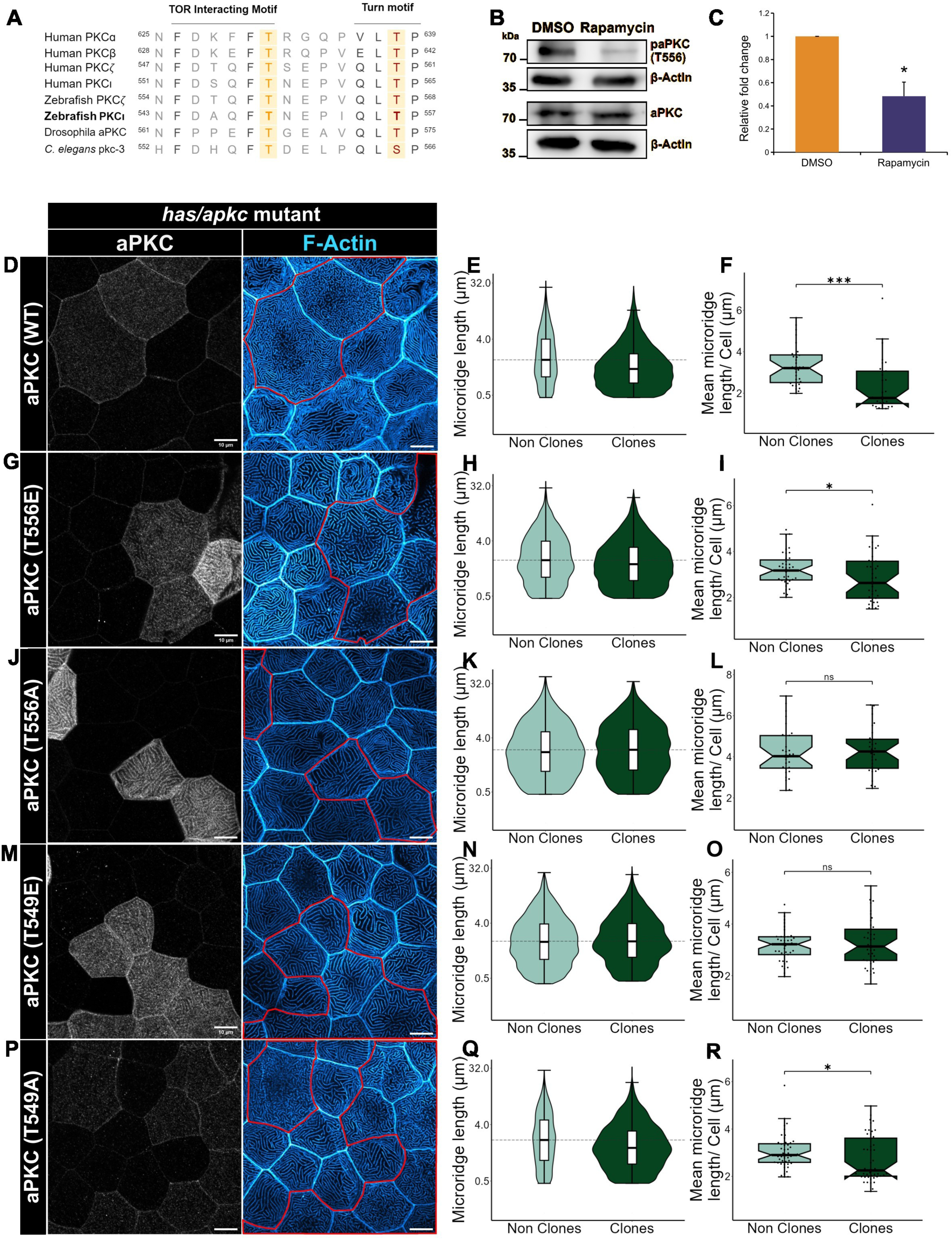
mTORC2 controls microridge elongation via phosphorylation of aPKC at TIM and turn motif. (A) Sequence homology alignment of TIM and turn motif of Human, Zebrafish, Drosophila and *C.elegans* PKCs including both conventional and atypical PKCs. Highlighted Threonine residues are potential mTORC2 phosphorylation sites in zebrafish and are relevant for this study. Western blot analysis (B) and quantification (C), using anti phospho aPKC Thr 555 antibody (Thr 556 in zebrafish), of the phosphorylation level at Thr 556 of aPKC in zebrafish embryos treated with DMSO and rapamycin for 48 hrs. Note that the levels of Phospho-aPKC and aPKC were normalized to their corresponding loading control (β-Actin). (D) Representative images of peridermal cells of rhodamine phalloidin stained *has/apkc* (*prkci^-/-^*) embryos at 48 hpf (D, G, J, M, and P) and distribution of microridge lengths (E, H, K, N, and Q) and mean microridge lengths (F, I, L, O, and R) from these embryos injected with plasmids containing HA-tagged aPKC wild type (D-F), aPKC (T556E) (G-I), aPKC (T556A) (J-L), aPKC (T549E) (M-O) and aPKC (T549A) (P-R). Note that embryos were immunostained with aPKC antibody (Grey in D, G, J, M, and P) for identifying clones. Experiments represented in (D-F), (G-I), and (P-R) were performed four times and the rest were performed three times, independently. For data represented in (E, F) n = 32 cells from N = 10 embryos, (H, I) n = 39 from N =15, (K, L) n = 31 from N = 7, (N, O) n = 41 from N = 9, and (Q, R) n = 48 from N = 10, were analyzed. Asterisks represent the significant difference by Mann Whitney U test for (F, I, L, O, and R) and student’s t-test for (C). Error bars in (C) denote SEM. *ns* (non significant) p>0.05, * p<0.05, *** p<0.001. The scale bar corresponds to 10μm.

To test the significance of this phosphorylation in microridge patterning, we generated phospho-mimetic (aPKC T556E) and phospho-defective (aPKC T556A) mutant variants of zf-aPKC. To assess the function of these mutant aPKC variants, we clonally expressed them in *has/apkc* mutants by injecting plasmids at one cell stage and analyzed the microridge phenotypes at 48 hpf. We reasoned that the functional forms of aPKC will rescue the long microridge phenotype displayed by the *has/apkc* mutant embryos. Consistent with this, clonal expression of wild-type aPKC resulted in the rescue of the long microridge phenotype observed in *has/apkc* mutant peridermal cells compared to the surrounding non-clonal cells, suggesting that the assay works well (fig. 4D-F). Clonal expression of the phospho-mimetic turn motif variant of aPKC (aPKC T556E) in the *has/apkc* mutant peridermal cells showed a significant rescue in the microridge phenotype (fig. 4G-I). On the other hand, a phospho-defective mutant of the same residue (aPKC T556A) did not show rescue in the microridge phenotype in the clonal cells (fig. 4J-L). These results suggest that the turn motif phosphorylation by mTORC2 is important for aPKC activity and thus regulation of the microridge length.

Recently, another threonine residue juxtaposed to the turn motif Thr in PKC family proteins has been proposed as a potential site for mTORC2 phosphorylation, which was termed the ‘TOR interaction motif’ or TIM (Baffi et al., 2021). In PKCβII, mTORC2-mediated phosphorylation at both TIM and turn motifs is required for the monomerization of the native PKC protein which is needed for its priming (Baffi et al., 2021). In aPKCλ/ɩ, this threonine site is also conserved, and it is adjacent to the turn motif threonine (fig. 4A, supp fig. 3F) raising a strong possibility of it being functionally relevant. Therefore, we investigated any potential role of TIM phosphorylation in aPKC activity. As before, we created phospho-mimetic (aPKC T549E) and phospho-defective (aPKC T549A) mutant variants of aPKC to genetically assess the functional significance of the TIM site. The clonal expression of the phospho-mimetic version of the aPKC at the TIM site (aPKC T549E) did not rescue the microridge phenotype in the clonal cells of *has/apkc* periderm (fig. 4M-O). On the other hand, the phospho-defective version (aPKC T549A), showed a significant rescue in the microridge length when clonally expressed compared to the surrounding cells (fig. 4P-R). These data indicate that phosphorylation, if any, at the TIM site would negatively regulate aPKC activity *in vivo*.

Further, we used a pan-phosphothreonine antibody to test whether the TIM Thr 549 is indeed phosphorylated *in vivo*. We expressed HA-tagged aPKC wildtype and the phospho-defective (T549A, T556A, and T549A/556A) variants in HEK293T cells and assessed the aPKC phosphorylation using the pan-phosphothreonine antibody. We reasoned that these mutations would inhibit Thr phosphorylations at the respective sites, decreasing the overall phosphorylation of the aPKC protein. The cells were lysed and immunoprecipitated using HA antibodies followed by western blot analysis. The quantifications revealed that the pThr level of the aPKC T556A version was significantly decreased compared to wildtype (supp fig. 3G, H). Importantly, T549 phospho-defective also showed a significant reduction in pThr level compared to the wildtype (supp fig. 3G, H) suggesting that the TIM gets phosphorylated *in vivo*. The double phospho-defective variant also showed a trend toward a combined reduction (supp fig. 3G, H).

To conclude, zebrafish aPKCλ/ɩ is phosphorylated at the threonine residue in the turn motif by mTORC2 and this phosphorylation is essential for its activity. On the other hand, the phosphorylation of threonine residue at the TIM site, presumably via mTORC2, is detrimental to aPKC activity.

### Differential phosphorylation of aPKC at TIM and Turn motif is essential for its activity

A reduction in aPKC activity upon TIM phospho-mimetic modification was an interesting finding. Given the adjacent position of the turn motif (supp fig. 3F) with respect to TIM, we asked whether phosphorylation status at Thr549 of TIM influences the functional outcomes achieved via the turn motif phosphorylation. To test this, we combined both the phospho-mimetic mutations creating aPKC T549E/556E variant and clonally expressed in *has/apkc* mutant embryos as before. We did not observe any rescue in microridge length phenotype in the clones expressing the double phospho-mimetic variant (fig. 5A-C) indicating this variant was non-functional. On the other hand, a double phospho-defective (aPKC T549A/556A) aPKC variant also did not show rescue, validating the importance of the turn motif phosphorylation (fig. 5D-F). Effectively, these results suggest that suppressive TIM phosphorylation influences turn motif phosphorylation and the simultaneous phosphorylation or dephosphorylation at both the sites of aPKC downregulates its functional activity.

**Figure 5.**
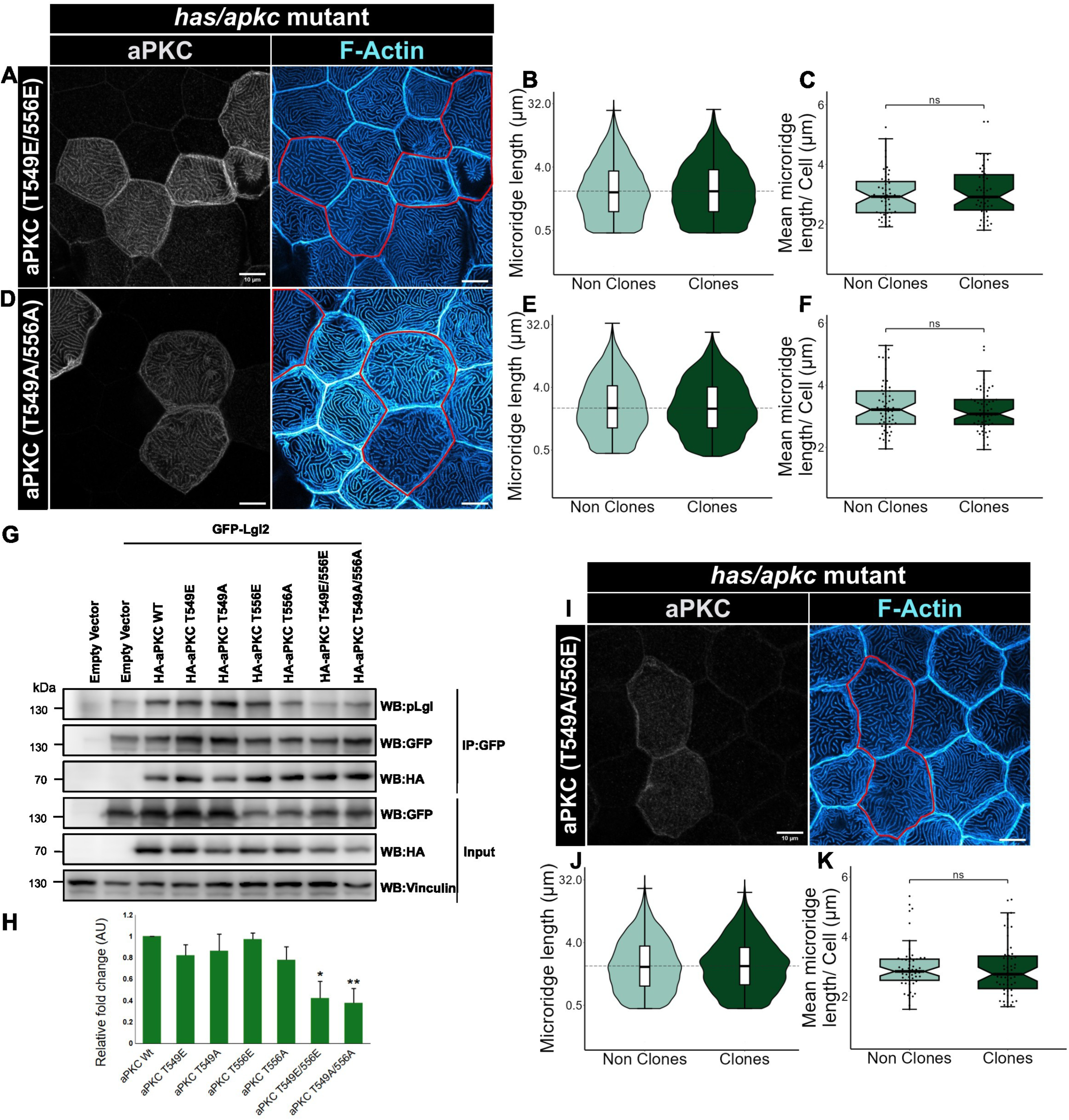
Simultaneous phospho-mimetic or phospho-defective state at both TIM and turn motif inhibits aPKC activity. Confocal images of peridermal cells from aPKC and rhodamine-phalloidin stained *has/apkc* (*prkci^-/-^*) embryos at 48 hpf (A, D, and I), and graphical representation of the distribution of microridge lengths (B, E, and J), and mean microridge lengths (C, F, and K) from these embryos injected with HA tagged aPKC (T549E/556E) (A-C), aPKC (T549A/556A) (D-F) or aPKC (T549A/556E) (I-K) plasmids. (G) Western blot shows changes in the level of pLgl2 (Ser 645/649) upon co-expressing the Lgl2 plasmid with different variants (as denoted) of aPKC in HEK293T cells and pulling down using GFP antibody. The relative difference in the pLgl2 level is quantified and represented in (H). Experiments represented in (I, K) were performed four times; the rest were performed three times independently. For data presented in (B, C) n = 50 cells from N = 15 embryos, (E, F) n = 61 from N =13, (J, K) n = 54 from N = 16 were analyzed. Asterisks denote significant differences when analyzed using the Mann-Whitney U test in (C, F, and K) and using the student’s t-test in (H). The error bars in (H) denote SEM. *ns* (non significant) p>0.05, * p<0.05. The scale bar corresponds to 10μm.

To assess the aPKC kinase activity of various phospho-mimetic or phospho-defective versions, we performed an *in vivo* kinase assay. As before, we expressed aPKC mutant variants in HEK293T cells along with its substrate Lgl2 tagged to GFP. One day after transfecting with the aPKC and Lgl2 plasmids, we isolated cell lysate and pulled down Lgl2 using an anti-GFP antibody to probe for its phosphorylation at Ser 645/649 (643/647 in zebrafish) which is known to be regulated by aPKC (Betschinger et al., 2003). Consistent with the genetic analysis, we observed no change in aPKC activity in turn motif phospho-mimetic variant (aPKC T556E) (fig. 5G, H) as compared to the wild-type aPKC. There was a marginal reduction in the aPKC activity in TIM phospho-mimetic (aPKC T549E) as well as turn motif phospho-defective (aPKC T556A) variants (fig. 5G, H), which seems consequential given that the genetic data suggested that these modifications inhibit the aPKC function. Importantly, there was a significant decrease in the aPKC activity when double mimetic or defective modifications were present, suggesting that simultaneous phosphorylation or dephosphorylation at both sites (T556 and T549) reduced the kinase activity of aPKC (fig. 5G, H).

Next, we asked whether combining the TIM phospho-defective mutation with the turn motif phospho-mimetic (aPKC T549A/556E) would yield a synergistic gain in the aPKC activity given these single mutant variants could significantly rescue the microridge phenotype in *has/apkc* mutant embryos. Surprisingly, aPKC T549A/556E was unable to rescue microridge length phenotype in *has/apkc* mutant peridermal cells (fig. 5I-K) suggesting that dynamic and differential phosphorylation of these two residues must be required to make aPKC fully functional.

To further demonstrate that the phosphorylation status at TIM and turn motif directly affect aPKC kinase activity, we used a constitutively active (CA) mutation of aPKC. As previously reported, aPKC activity is blocked by its pseudo-substrate (PS) region in the absence of any substrate. Introducing a positively charged amino acid in this PS region leads to constitutive activation of aPKC (Graybill et al., 2012). We mutated the conserved alanine 122 of PS to glutamate in aPKCλ/ɩ and expressed it in *has/apkc* mutants as before. As anticipated, we observed a significant rescue in microridge length in the clones expressing CA-aPKC (A122E) (fig. 6A-C). When we combined the A122E mutation with inhibitory mutations of aPKC - i.e. turn motif phospho-defective (aPKC A122E/T556A) or TIM phospho-mimetic (aPKC A122E/T549E) - we observed that CA mutation could significantly rescue microridge length in both the conditions (fig. 6D-F and 6G-I, respectively). However, the CA mutation was unable to rescue the activity of aPKC with double phospho-mimetic or phospho-defective mutations (fig. 6J-O). These data suggest that simultaneous phosphorylation or dephosphorylation states may directly reduce the kinase property of aPKC or affect substrate interaction. However, we ruled out the latter possibility of effect on the substrate interaction given that the overall interaction of Lgl2 with aPKC was not seen to be reduced in the *in vivo* kinase assay (fig. 5G).

**Figure 6.**
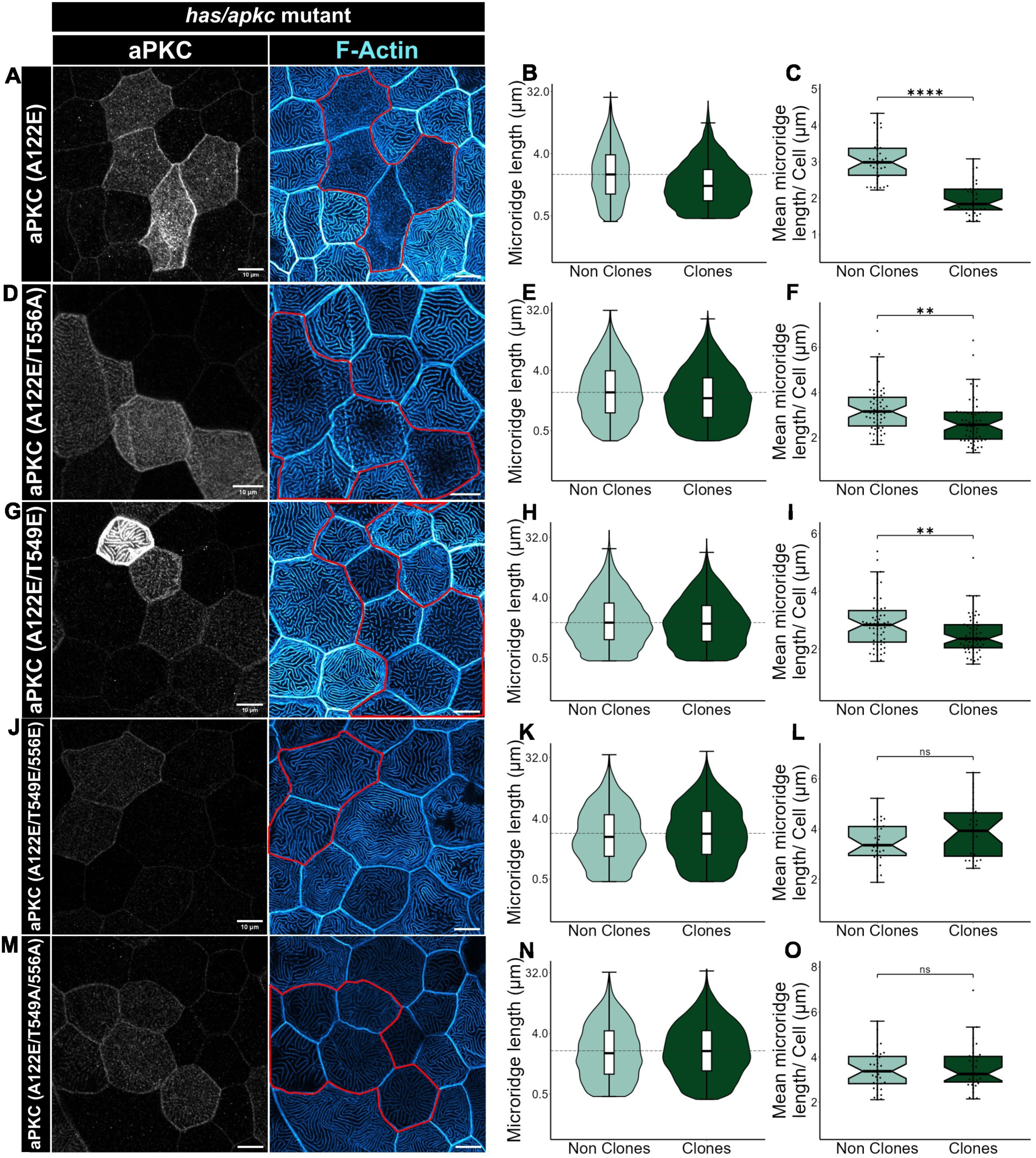
Constitutively active aPKC does not rescue the effect of simultaneous phospho-modifications at TIM and turn motif. Representative images of peridermal cells from aPKC and rhodamine phalloidin stained *has/apkc* (*prkci^-/-^*) embryos at 48hpf (A, D, G, J, and M) and violin plots of microridge length distributions (B, E, H, K, and N) and box-jitter plots of mean microridge lengths (C, F, I, L, and O) for these embryos injected with HA-tagged constitutively active aPKC (A122E) (A-C), aPKC (A122E/ T556A) (D-F), aPKC (A122E / T549E) (G-I), aPKC (A122E/T549E/556E) (J-L) and aPKC (A122E/T549A/556A) (M-O) plasmids. All the experiments were performed three times independently. For data presented in (B, C) n = 38 cells from N = 10 embryos, (E, F) n = 59 from N = 16, (H, I) n = 59 from N = 16, (K, L) n = 31 from N = 10, and (N, O) n = 35 from N = 10 were analyzed. Asterisks denote the significant difference when the data were analyzed using the Mann-Whitney U test for all the box-jitter plots. *ns* (non significant) p>0.05, ** p<0.01, **** p<0.0001. The scale bar corresponds to 10μm.

In conclusion, mTORC2 positively regulates aPKC when it phosphorylates the turn motif, which is crucial for microridge patterning. However, phosphorylation at the TIM site results in the downregulation of aPKC activity; it also masks the effect of turn motif phosphorylation. Of note, the persistent dephosphorylation status at the TIM site in combination with turn motif phosphorylation is also detrimental to aPKC’s activity suggesting the requirement of transient phosphorylation at the TIM site for the activity.

### TIM motif phosphorylation leads to decreased interaction between pThr556 of the turn motif and Arg254 of the glycine-rich motif

Our *in vivo* kinase and genetic assay both show that the incorporation of simultaneous phospho-mimetic mutation at the TIM (T549E) site and the turn motif (T556E) suppresses aPKC activity, whereas the single mutant T556E is active. We sought to determine the reason for this inhibition of kinase activity for the double mutant by using in-silico methods. We first checked whether the mimetic mutation has any effect on the protein structure itself. We used a recently developed method of analyzing the effect of amino acid mutations using effective strain (ES) as a metric (McBride et al., 2023). The effective strain measures the structural deformation in proteins due to a mutation and was employed by McBride et al. to show that AlphaFold2 generated structures were sensitive to single amino acid mutations. We generated the 3D AlphaFold2-predicted structures of wildtype and different mutant variants of aPKC protein using the ColabFold server (Mirdita et al., 2022). We then calculated the relative effective strain in either single or double mutants versus wildtype structures (McBride et al., 2023). Our analysis shows that there is an increase in ES at the TIM site in both aPKC T549E and aPKC T549E/556E (fig. 7A, C) as compared to wild-type aPKC; this increase was not observed in the turn motif phospho-mimetic version (T556E) (fig 7B, C). In contrast, there is not much change in ES at TIM or turn motif of phospho-defective versions (aPKC T549A, aPKC T556A, or aPKC T549A/556A) relative to wild-type (fig. 7 A-C). Moreover, in the double phospho-mimetic version (aPKC T549E/556E), the relative ES for the turn motif is higher (fig. 7C, D) compared to aPKC T556E. These data indicate that the phosphorylation at T549 induces a strong local conformational strain as well as a weaker non-local strain at the turn motif in the protein.

**Figure 7.**
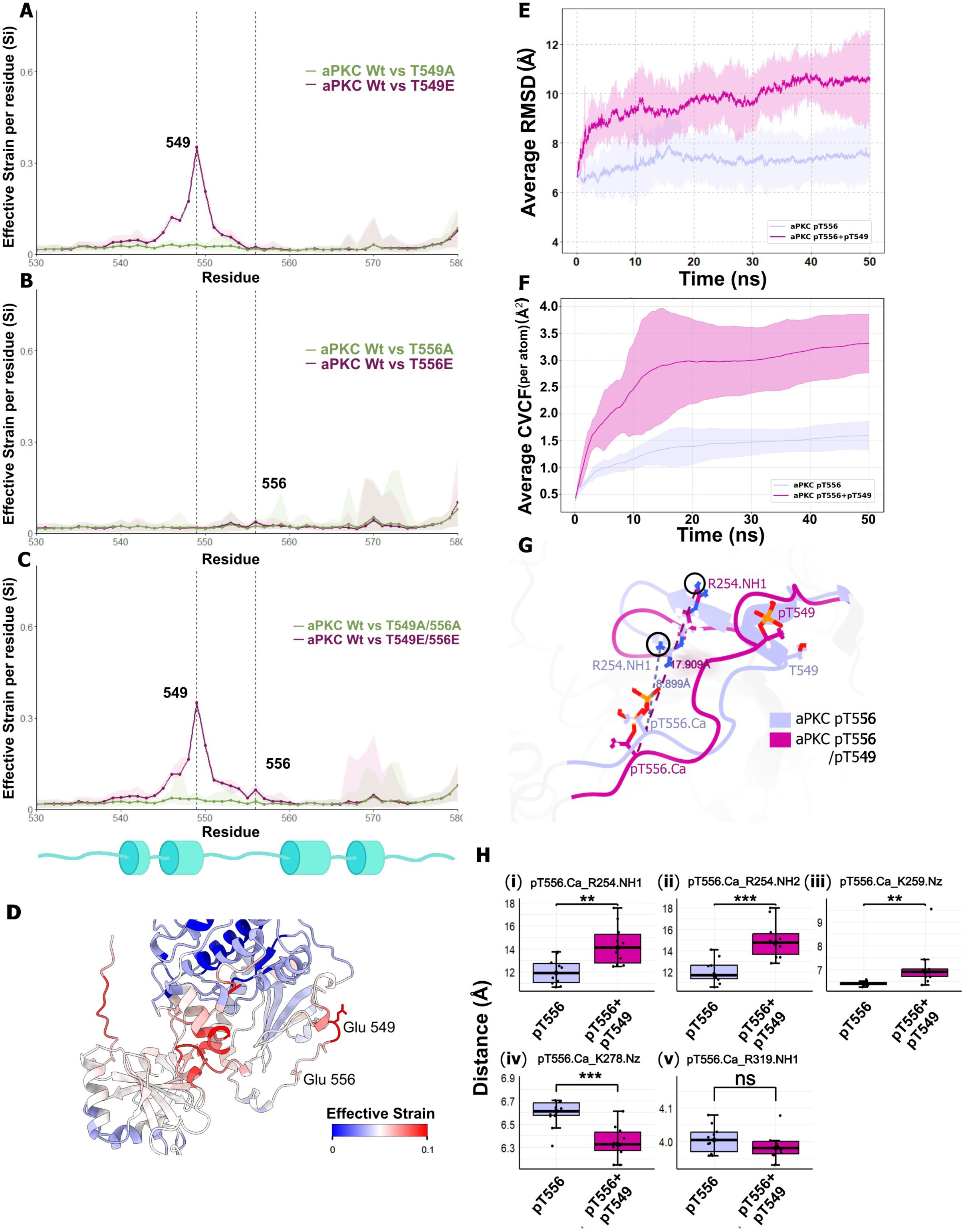
TIM phosphorylation increases local strain and distance between pT556 and R254 of the glycine-rich motif. Line plot for effective strain (Si) calculated per residue between (A) aPKC Wt vs aPKC T549E (Magenta) and aPKC Wt vs aPKC T549A (Green), (B) aPKC Wt vs aPKC T556E (Magenta) and aPKC Wt vs aPKC T556A (Green), and (C) aPKC Wt vs aPKC T549E/556E (Magenta) and aPKC Wt vs aPKC T549A/556A (Green). Note that Si was calculated from five top AF2 predictions of the 3D model compared against each other. The average trend line (darker) with min and max Si (lighter shade) is represented in the plot. For better visualization, only residues from 530-580 are shown. (D) Visual representation of Si per residue of aPKC T549E/556E. The heat map denotes an index of low to high effective strain. The Average RMSD (E) and CVCF (F) of all backbone atoms calculated after simulating phosphorylation at either T556 (purple) or both at T556 and T549 (magenta) of aPKC. The shaded region denotes standard deviation across all the trajactories generated from MD simulation. (G) Visual comparison of distances between pT556.Cɑ of turn motif and R254.NH1 of Glycine-rich motif of simulated aPKC pT556 (purple) and aPKC pT556/pT549 (magenta). (H) Box-jitter plots (i)-(v) comparing the average distances between the residues mentioned in Supplementary table 1 among aPKC pT556 and aPKC pT556/pT549 obtained from 10 iterations of MD simulation. Asterisks denote the significant difference by the student’s t-test for all box-jitter plots. *ns* (non significant) p>0.05, * p<0.05, ** p<0.01, *** p<0.001.

To further consolidate our hypothesis, we carried out an all-atom Molecular Dynamics (MD) Simulation study of the differential effect of single versus double phosphorylations on aPKC structure. Specifically, we added a phosphate group at T556 or both at T549 and 556 in aPKC wildtype structure and monitored changes in the key inter-residue separations. Our analysis was based on 10×50 ns NPT production runs of the two aPKC systems each solvated with water and K^+^ and Cl^-^ ions. The average RMSD of the backbone atoms relative to the starting equilibrated structure in the trajectories (fig. 7E, supp fig. 4A, B) stabilizes earlier (∼20 ns) as compared to the double mutant (∼35 ns). Furthermore, the average RMSD with respect to the starting reference structure is higher for the double phosphorylated aPKC (fig. 7E) indicating an ∼3Å greater drift as compared to pT556. Next, we analyzed the cumulative variance of coordinate fluctuations (CVCF) for all-atoms of the two aPKC phosphorylated variants, which provides a more nuanced view of the underlying energy landscape (Paul et al., 2020). Essentially, plateaus in the CVCF trace of a single protein trajectory indicate local equilibria with Boltzmann population statistics. The average CVCF values extracted at plateaus and standard deviations derived from multiple protein trajectories provide an estimate of the effective curvature and roughness of the energy landscape, respectively. For the aPKC systems, the per-atom CVCF traces for structured backbone atoms (Supplementary table 2) reveals that the energy landscape of pT556+pT549 double mutant is more rugged (larger standard deviations) than that of the pT556 single mutant (fig. 7F). The average per atom CVCF traces across the full set of MD trajectories do not show plateaus for either system, indicating that both single and double mutant systems have yet not achieved equilibration (across all trajectories) at 50 ns (fig. 7F). However, the slopes are higher for the double mutant indicating that this system is more unstable relative to the single mutant. Furthermore, the higher per-atom CVCF of aPKC pT556+pT549 (fig. 7F) suggests an additional phosphorylation at T549 increases flexibility of aPKC compared to only T556 phosphorylation.

Although the overall average per-atom CVCF trace across all MD trajectories for both systems exhibits significant drifts, individual trajectories display a larger number of plateaus signifying local equilibration (supp fig. 4C). Our analysis shows that aPKC pT556 has CVCF trace segments with longer plateaus compared to aPKC pT556+pT549 (supp fig. 4C) suggesting a greater number of stable states which allow for establishing local equilibria. We further quantified the stability of accessible states by calculating the average residence time of the plateaus across all trajectories for both systems. The results indicate that aPKC pT556 exhibits more stable states, as evidenced by its higher average residence time for each threshold (supp fig.4D).

A previous study analyzed the crystal structure of human aPKC and described the formation of the ATP-binding pocket in PKCɩ (Takimura et al., 2010). It was shown that the phosphate group of the turn motif threonine forms stable interactions with the kinase core residues (Supplementary table 1) which is necessary for ATP binding and phosphate transfer. This led us to investigate whether adding a phosphate group at T549 affects this interaction. We calculated the distance between Cɑ of Thr556 and those of the kinase core residues mentioned in Supplementary table 1 at the relative equilibrated state. We found that the average distance between the Cɑ atoms of pT556 and the R254 and K259 of the Glycine-rich motif significantly increased (fig. 7G and 7H(i)-7H(iii)) in the aPKC when a phosphate group was present at T549 residue (T556_Cɑ_-R254_NH1_ pT556:∼12.03 Å, pT556+pT549:∼14.34 Å; T556_Cɑ_-R254_NH2_ pT556:∼12.05 Å, pT556+pT549:∼15 Å; T556_Cɑ_-K259_Nz_ pT556:∼6.04 Å, pT556+pT549:∼7.1 Å). This is interesting as the glycine-rich motif is important in ATP binding (Hemmer et al., 1997).

We next investigated the differences in active site dynamics by applying the CVCF-trace analysis for all atoms belonging to residues within 3-10 Å of ATP (supp fig. 4E-H). We find that while the active site dynamics in the immediate vicinity (within 3 Å) of the ATP for the aPKC pT556+pT549 and aPKC pT556 variant are comparable at 50 ns, the dynamics for the two variants is distinct at larger cutoffs (supp fig. 4E-H). Specifically, the extended active site comprising of residues 5 Å or more away from the ATP is significantly more dynamic for the double phosphorylated aPKC relative to that in aPKC pT556.

In conclusion, phosphorylation at TIM in aPKC leads to structural deformation leading to increased distance between pThr556 and glycine-rich motif which could potentially affect the ATP binding site. Furthermore, double phosphorylation leads to a more unstable protein fold with significantly higher flexibility compared to single phosphorylation at pThr556.

## Discussion

Apical projections aid in epithelial functions such as absorption, secretion, diffusion, signaling, etc. Their formation is regulated by cell-autonomous mechanisms that control cell polarity, intracellular transport and the formation and regulation of F-actin. Here we identify mTOR as a signaling pathway that acts upstream of both actin and polarity regulators to control microridges, F-actin-based apical projections present on squamous epithelial cells. Our analysis shows that mTORC1 directly influences the microridge formation via the actin cytoskeleton regulators. By downregulating the activity of RhoA via S6K1, mTORC1 prevents overactivation of NMII plausibly by ROCK-mediated phosphorylation (fig. 8). Indeed, NMII is known to orchestrate the development of microridges by regulating dynamic fission-fusion events via local actomyosin contraction as well as apical constriction (Bhavna & Sonawane, 2024; Raman et al., 2016; van Loon et al., 2020, 2021). On the other hand, mTORC2 regulates activity of aPKC by its differential phosphorylation at TIM and turn motifs. aPKC restricts the microridge elongation via controlling the localisation of Lgl, which in conjunction with NMII, functions in microridge elongation (Raman et al., 2016; fig. 8). Amongst other apical projections, functions of mTORC1 or mTORC2 are also essential for maintaining microvilli length in the proximal tubule cells (Grahammer et al., 2017). Furthermore, the regulation of primary cilia growth by mTORC1 is likely to be context-dependent (Sherpa et al., 2016; Takahashi et al., 2018). It will be interesting to test whether mTOR also regulates these other projections via polarity proteins or actin regulators.

**Figure 8.**
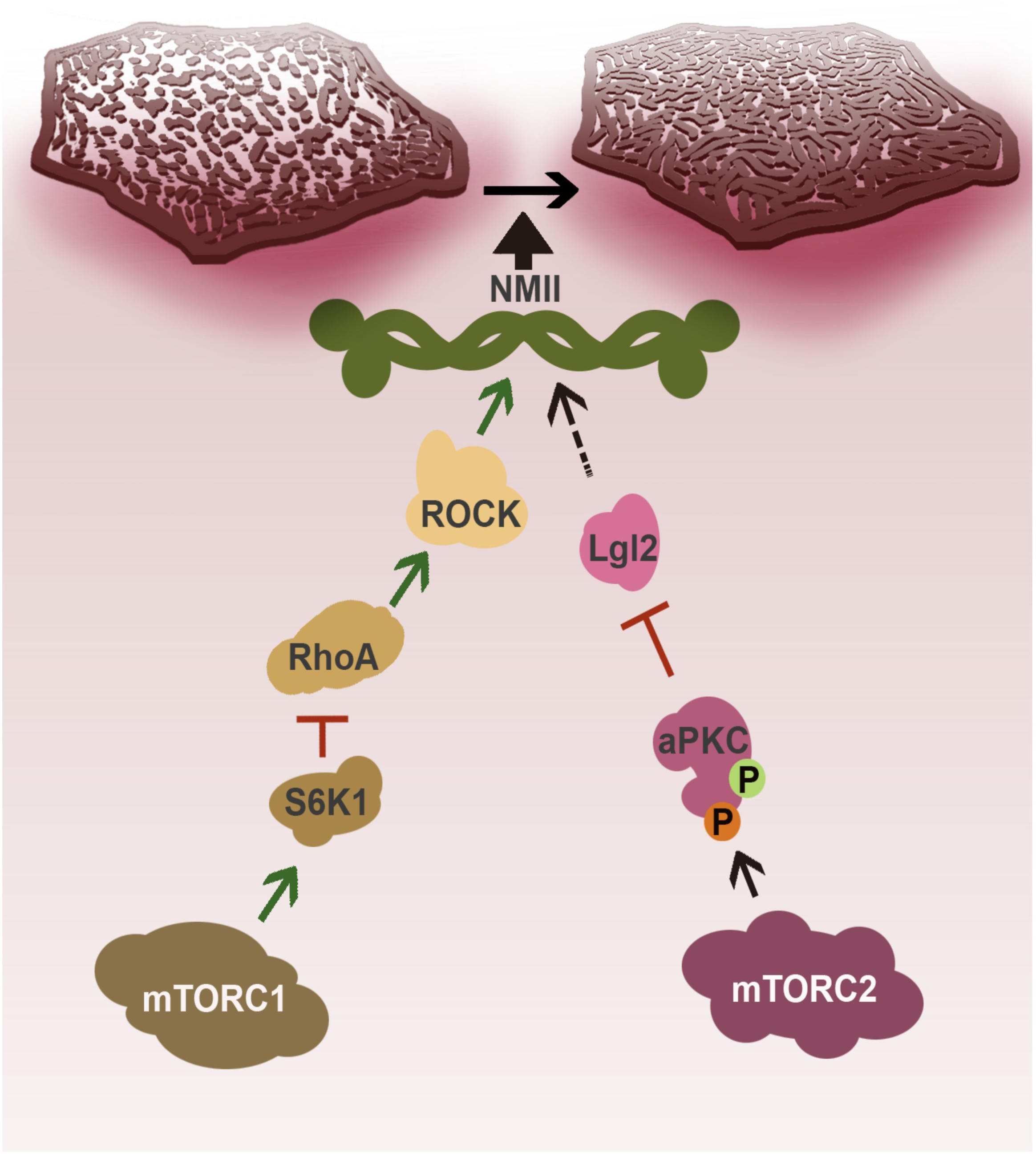
Schematic of microridge pattern regulation by mTORC1/S6K1 and mTORC2/aPKC axes. Proposed model representing the two mTOR signaling axes regulating the NMII activity. mTORC1 and its substrate S6K1 downregulates the RhoA/ROCK activity whereas, mTORC2 regulates aPKC activity via differential phosphorylation. Both of these axes converge on NMII which influences microridge pattern formation.

mTORC1 is primarily a central regulator of cell metabolism, growth, and proliferation (Panwar et al., 2023). However, its role in the regulation of actin cytoskeleton is not clear. A few studies have shown that mTORC1 regulates the gene expression or protein translation of actin regulators such as RhoGTPases and Arp2/3 (Liu, Luo, et al., 2010; Zhao et al., 2021). In this study, we report an unconventional role of mTORC1 in the regulation of microridges by restricting RhoA activity via S6K1-mediated inhibition. The mTORC1/S6K1 axis, which links the growth factor or nutrient inputs with cellular physiology, is also important in microridge pattern development. In this regard, how S6K1 inhibits RhoA activity needs to be investigated in the future. It was shown that phosphorylation of RhoA C-term leads to its sequestration by RhoGDIs preventing its membrane association (Ellerbroek et al., 2003). It is worth exploring whether similar phosphorylation is performed by S6K1 directly or via any kinase downstream of S6K1.

mTORC2 is known to regulate actin cytoskeleton via RhoA or Rac1 in migrating cells, neurons, and mesenchymal stem cells (Huang et al., 2013; Saha et al., 2023; Sen et al., 2014; Thompson et al., 2013). Independently, mTORC2 modulates the actin cytoskeleton via the regulation of PKC family proteins (Barandier et al., 2003; Matsuoka et al., 1998; Morrison Joly et al., 2017). mTORC2 also phosphorylates PKC at its turn motif and improves its stability (Baffi & Newton, 2022; Ikenoue et al., 2008; Li & Gao, 2014). Furthermore, mTORC2 activates PKCβII via its phosphorylation at the TIM site (Baffi et al., 2021; Baffi & Newton, 2022). However, it is unclear whether and how mTORC2 regulates aPKC in an *in vivo* context. Here we report that both these phosphorylation sites are conserved in zebrafish aPKCλ/ι and are phosphorylated. In fact, we show that the threonine at the turn motif is phosphorylated in an mTOR activity-dependent manner and this phosphorylation of aPKC is important for its activity and microridge pattern regulation. Intriguingly, the phosphorylation at the TIM motif of aPKC is detrimental to its activity endowed via the turn motif phosphorylation. Our MD simulation data suggests that the phosphorylation at the threonine of the TIM motif decreases the protein stability. In addition, this phosphorylation increases the distance between the turn motif and the arginine group of the glycine-rich motif of aPKC (fig 7G, H). As previously reported, the Glycine-rich motif is crucial for ATP binding and stability (Hemmer et al., 1997). Additionally, we also observe a destabilizing effect of TIM phosphorylation on the overall protein fold relative to a single phosphorylation at the turn motif. These MD simulation analyses corroborate our genetic and biochemical data, which show the reduced activity of doubly phospho-mimetic aPKC. Based on the MD data, we propose that TIM phosphorylation affects aPKC functions by reducing its overall stability which may possibly impact ATP binding. This aspect needs to be verified further with explicit ATP binding analysis and experiments.

How does this dual phosphorylation regulate the activity of zebrafish aPKCλ/ι? TIM and turn motif phosphorylation by mTORC2 in a dimerized native PKCβII leads to its monomerization, which is followed by a series of phosphorylations priming it for further interaction with its substrate (Baffi et al., 2021). It has been reported that native aPKC does not dimerize (Le et al., 2022). Hence, it is likely that the dual phosphorylation by mTORC2 affects aPKC differently than PKCβII. It has been shown that turn motif phosphorylation is co-translational in aPKC (Tobias et al., 2016). We hypothesize that TIM phosphorylation prevents aPKC activation at non-apical localization since the turn motif is co-translationally phosphorylated. However, persistent phosphorylation at TIM is also not desirable, as it suppresses aPKC kinase activity by reducing its stability and perhaps ATP binding. Only when aPKC localizes to the apical domain, the threonine at TIM may get dephosphorylated and regain its activity. This apically localized fully active aPKC can then prevent Lgl2 localization at the apical side by phosphorylation and thus control the microridge elongation (Betschinger et al., 2003; Raman et al., 2016).

In conclusion, our study provides a mechanistic understanding of mTORC1 and mTORC2 mediated microridge pattern regulation by converging on NMII (fig. 8). Signaling via mTORC1 and mTORC2 allows the cells to respond to a plethora of extrinsic cues such as growth factors, metabolic status, nutrient availability, and membrane tension (Panwar et al., 2023; Saha et al., 2023). We propose that in the developing squamous epithelium mTOR signaling integrates extrinsic cues, possibly including developmental or growth factor signals, with downstream cell-autonomous mechanisms to regulate tissue-wide formation of microridges.

## Materials and Methods

### Zebrafish Strains

The zebrafish husbandry and experiments were performed according to the guidelines of the institutional animal ethics committee approved vide TIFR/IAEC/2017-11 and TIFR/IAEC/2019-6. *Tübingen* wildtype (*Tü*) fishes and *heart and soul* (*has/apkc*) mutant strain (Horne-Badovinac et al., 2001) were used for the experiments mentioned in this article.

### Gene synthesis, cloning, site-directed mutagenesis, and microinjection of plasmids

N-terminal HA-tagged zebrafish aPKC wildtype CDS (NM 131855) synthesis in pCS2+8 vector (a gift from Amro Hamdoun; Addgene plasmid # 34931), and aPKC T556E and aPKC T556A site-directed mutagenesis (SDM) were done by GeneArt, Life Technologies. aPKC T549E, aPKC T549A, aPKC A122E, and the combined mutations were achieved by SDM with the primers mentioned in supplementary information 1, using Q5 Polymerase (Neb) according to the protocol published earlier (Edelheit et al., 2009). pcDNA3-EGFP-RhoA-Q63L (Addgene plasmid # 12968) was a gift from Gary Bokoch.

After isolation of plasmids, working solutions were made in nuclease-free water. About 45 pg of each plasmid was injected in one or two cell-staged embryos.

### Inhibitor treatments

Small molecule inhibitor treatments were administered in E3 buffer without methylene blue. For every condition, about 15–16 embryos at 24 hpf were taken in a six-well plate. DMSO as vehicle control and the inhibitors, with the final DMSO concentration of 1% in both cases, were added to the solution and incubated for 24 hours (unless specified otherwise) at 28.5 °C. Following this, three washes of five minutes each were given using E3 without methylene blue before fixing the embryos. Small molecule inhibitors used in this study and their final concentrations are as follows: Rapamycin (2μM) (SCBT, sc3504), Rhosin (40μM) (MedChemExpress, HY-12646), LY2584702 tosylate (10μM) (Sigma, SML2892), Blebbistatin (50μM) (Sigma, B0560), and Rockout (100μM) (SCBT, sc-203237).

### Protein sample preparation from embryos

For protein sample preparation about 25–30 embryos per condition were taken in a 1.5ml microcentrifuge tube. First, an E3 (without methylene blue) wash was given followed, by a wash with Ginsburg’s fish ringer solution without Ca^2+^ (111.2mM NaCl, 3.35mM KCl, 2.38mM NaHCO_3_). The embryos were incubated in fish ringer’s solution for 10 mins at 4 °C on a rotor. Using a 200 μl pipette tip, embryos were gently agitated to remove the yolk, following which they were kept at 16 °C for 10 mins on a thermomixer at 1200rpm. Then the solution was discarded, and the embryos were given three washes of PBS (137mM NaCl, 2.7mM KCl, 10mM Na_2_HPO_4_, 1.8mM KH_2_PO_4_, pH 7.4). After removing PBS completely, 60-80 μl RIPA Lysis buffer (50 mM Tris-Cl; pH 7.4, 150 mM NaCl, 0.1% SDS, 0.5% Na-deoxycholate, 1% NP-40, and 0.01% Sodium azide) including 1X PhosStop (Roche, PHOSS-RO), 1mM NaF, 1X PIC (Sigma, S8830), and 1X PMSF was added. The embryos were sonicated at 20% amplitude for 50 secs with 10 sec on and off pulses. The tubes were microcentrifuged for 30 mins at 13,000 rpm at 4 °C and the supernatant was collected. Finally, samples were added with Laemmlie buffer, boiled, and stored at -20 °C until further use for SDS-PAGE electrophoresis.

### RNA isolation, cDNA synthesis, and qPCR assay

For isolating RNA, about 30 embryos were taken in a 1.5 ml microcentrifuge tube. After removing the excess E3 buffer, 200 μl chilled TRIzol (Invitrogen, 15596026) was added. The embryos were crushed carefully with a plastic pestle while keeping the tube in ice. Following this, 800 μl additional TRIzol and 200 μl of chilled chloroform were added to the tube. After mixing and centrifugation, the RNA in aqueous phase was collected in a fresh tube and 1ml of 100% Isopropanol was added for precipitation. During precipitation,1 μl GlycoBlue (Invitrogen, AM9516) was added, and the sample was left at -20 °C overnight. The RNA pellet obtained after centrifugation was washed with 75% Ethanol followed by drying at 45 °C. Finally, the RNA was dissolved in nuclease-free water.

To make cDNA, SuperScript IV First-Strand synthesis kit (Invitrogen, 18091050) was used and instructions from the manufacturer’s manual was followed. For performing qPCR, we used the GoTaq qPCR master mix (Promega, A6001). The primers used for the qPCR are mentioned in supplementary information 1. For all the conditions, the qPCR was done in triplicates and later the Ct values were averaged. The expression of the gene of interest was determined after normalizing against the control gene *eef1a1*.

### Immunostaining

For immunostaining, embryos were first fixed with 4% PFA in PBS for either half an hour at room temperature, followed by overnight at 4 °C or four hours at room temperature. After fixation, the embryos were given two 5-minute PBS washes and permeabilized with PBT (0.8% Triton X-100 in PBS) by giving five 10-minute washes. 10% NGS (Jackson Immuno Research Labs, 005-000-121) in PBT solution was used as a blocking reagent for 3 hours at room temperature with a constant rotation on a slanted rotor. Primary antibodies were made in PBT with 1% NGS and the embryos were incubated overnight at 4 °C on a rotor. After the incubation, five PBT washes were given for half an hour each. The secondary antibody incubation was done on a rotor for 3 hours at room temperature. After giving seven 10-minute PBT washes, the embryos were post-fixed in PFA for 30 minutes at room temperature. Finally, after two PBS washes, the embryos were serially upgraded using 30%, 50%, and 70% Glycerol and kept at 4 °C until imaged.

For pAkt staining, embryos were fixed using freshly prepared chilled EAF (40% Ethanol, 5% Acetic acid, and 4% Formaldehyde in PBS) overnight. For immunostaining, the same protocol was followed as mentioned, with the addition of an antigen retrieval step which was done after permeabilization. For the antigen retrieval, the embryos were first equilibrated with 10mM Tris-Cl (pH-9) and 1mM EDTA solution for 10 mins on a rocker at room temperature, followed by an hour at 60 °C on a dry bath. After cooling down the solution, one PBT wash for 10 mins was given, and then proceeded with the blocking step. The embryos were incubated with pAkt antibody for a minimum of 16 hours. For phalloidin-only staining, the embryos were fixed with PFA for 2.5 hours at room temperature. After two PBS and four PBT washes, embryos were directly incubated with phalloidin solution for 2 hours at room temperature. Then the embryos were washed with PBT 6 times for 10 mins each and fixed again with PFA for half an hour followed by glycerol upgrade. The primary antibodies and the dilutions used were aPKC (SCBT, sc-17781, 1:750), pS6 (S240/244) (CST, 2215S, 1:100), pAkt (S473) (CST, 4060S, 1:50), pMLC (S19) (CST, 3671, 1:25), Lgl2 (Sonawane et al., 2009, 1:400), Ecad (BD Bioscience, 610182, 1:200), GFP (Torrey Pines, TP401 1:200), HA (Roche, 11867423001, 1:750). The secondary antibodies used were Alexa Fluor 488 (Invitrogen, A11029/A11034, 1:250), Cy3 (Jackson Immuno research lab, 115-165-146/111-165-144, 1:750), Cy5 (Jackson Immuno research lab, 115-175-146/111-175-144, 1:750), phalloidin 488 (Invitrogen, A12379, 1:200 or 1:400), phalloidin rhodamine (Invitrogen, R415, 1:200 or 1:400).

### Image acquisition and analysis

For confocal imaging, the embryos were mounted as mentioned before (Gupta & Sonawane, 2020). The confocal images were acquired on Zeiss 880 using 40X or 63X Plan-Apochromatic Oil immersion objectives at 2X and 1.3 or 1.5X optical zoom, respectively. The 12 or 16 bit images were taken at 1024X1024 pixels with 4 times line averaging. The parameters were kept constant during image acquisition except for the clonal images, for which the parameters were adjusted according to the expression level to maintain the dynamic range. All the images were analyzed and quantified using Fiji (Schindelin et al., 2012).

### Microridge quantification

Microridge lengths were quantified on ImageJ using a macro. Briefly, the brightness and contrast of the image were adjusted so that the ridges were properly visible. The image was smoothened twice. ‘Subtract Background’ was used with a rolling ball 500 to reduce the background noise. The image was then converted to 8 bit and ‘Auto Local Threshold’ was used with the Otsu method keeping the radius at 8 and other default parameters. Then the quantitative features were extracted using an algorithm written in ImageJ with an adaptation from a code by Ron DeSpain 2011 (DeSpain & Niccoli, 2011). The full version of the code can be found in the supplementary information 2. In comparison to the previous methods (Raman et al., 2016), this new algorithm fragments the microridges at the branch points, when present, to yield lengths of the unbranched microridge segments and unbranched microridges.

### Intensity quantification

Protein intensity was measured using Fiji (Schindelin et al., 2012). For pS6 intensity quantification in the periderm, Ecad or phalloidin was used as a cell boundary marker. First, a composite of pS6 and the other channel (Ecad or phalloidin) was taken. Next, from apical to basal slices, ROIs were drawn using the polygon tool and using the membrane marker as a reference. While staying on the pS6 channel, mean intensity was measured across the slices. Then an average intensity per cell was calculated and plotted. For pAkt or aPKC quantification, a segmented line tool with a width of 5 units was used to draw an ROI. This was done for all the slices where the membrane localization of the protein was visible. The mean intensity per slice was measured and the average per cell was calculated and plotted. For images taken at two different bit-depths (12 or 16), data with similar bit-depths were pooled and plotted, and the rest were plotted separately and presented in supplementary figure 1 A-C.

The intensity of western blots was quantified using Fiji. The box selector was used to select and measure the band intensity. The mean intensity of the bands was subtracted from background noise by selecting a nearby region. The mean intensity of the proteins was plotted after normalizing for their corresponding loading controls.

### Cell lines and reagents

HEK293T cells were grown in Dulbecco’s modified eagle medium (Gibco, Invitrogen, 11995-065) supplemented with 10% fetal bovine serum (MPBio, 092910154), and 100 U/mL Penicillin-100 μg/mL Streptomycin (Thermo Fisher Scientific, 15140122) at 37°C with a 5% CO2 in the environment. The cells were routinely tested for mycoplasma contamination.

### Cell transfection, immunoprecipitation, and western blot

Cells were transfected with the mentioned plasmids (fig. 5G and supp fig. 3G) using PEI Prime™ (Sigma, 919012). About 1.5-2 X 10^6^ cells were plated in 60 mm plates the day before transfection. The mentioned plasmids and PEI were mixed in Opti-MEM (Gibco, 31985062) separately and kept for 5 minutes at room temperature. Following this, the plasmid solutions and PEI were mixed in a ratio of 1:3 (plasmid:PEI) and kept for 20 mins at room temperature and added onto the cells. 2.5-3 μg HA-aPKC plasmids and 1.5 μg EGFP-xlgl2 (Chalmers et al., 2005) plasmids were used for the transfection. After 10 hours of transfection, media was changed to complete media for experiment presented in fig. 5G and to serum free media for experiment shown in supp fig. 3G. After overnight starvation the cells for experiment in supp fig. 3G were incubated with complete media for five hours before sample collection. 24 hours post-transfection, cells were lysed using non-denaturing lysis buffer (50mM Tris-Cl; pH 7.5, 150mM NaCl, 1mM EDTA, 10% Glycerol and 0.9% NP-40 including 1X PhosStop, 1mM NaF, 1X PIC, and 1X PMSF). For immunoprecipitation, protein A/G beads (Sigma, LSKMAGAG02) were incubated with lysis buffer for 3–4 hours before use. 20 μl beads were added to the cell lysate and incubated on an end-to-end rotor for 2.5-3 hours for pre-clearing at 4 °C. 1ug GFP antibody (Torrey Pines, TP401 1:200) for 1ug protein was added in the pre-cleared lysate and incubated on an end-to-end rotor overnight. Next, 40 μl of beads were added to the lysate and incubated for 3 hours at 4 °C. After the incubation, the beads were washed with the lysis buffer 3 times for 10 min each on a rotor at 4 °C. Finally, the beads were boiled in 2X Laemmlie buffer and the solution was collected. Protein samples were then electrophoresed on an SDS-PAGE. The proteins were transferred onto a PVDF membrane (Sigma, 03010040001) and blocked with 7% BSA (MPBio, 160069) for 2 hours at room temperature. Primary antibody incubation was done overnight at 4 °C and followed by three 10 mins TBST (0.1% Tween-20 in 1X TBS) washes. Secondary antibody incubation was done at room temperature for one hour followed by three 10-minute TBST washes. The membranes were developed using Supersignal ECL solution (Thermo Fisher A38554, 34579) and imaged using GE AI600. Primary antibodies used for western blot analyses are GFP (Torrey Pines, TP401, 1:2000), aPKC (Invitrogen, PA5-65216, 1:1000), pLgl (Abcam, 1:2000, ab59950* now discontinued), pThr (CST, 9381, 1:1000); HA (Roche, 11867423001, 1:1000), β-actin (Sigma, A1978, 1:2000), and Vinculin (Invitrogen, V4139, 1:1000). Secondary antibodies used are HRP-conjugated anti-mouse or anti-rabbit (Jackson Immuno research lab, 115-035-003/ 111-035-003, 1:30,000) and HRP-conjugated anti-rat (Sigma, A9037, 1:30,000).

### Molecular Dynamics Simulations

After obtaining Wild-Type aPKC structure using ColabFold (AlphaFold2 using MMseqs2) (Mirdita et al., 2022), post-translational modifications were introduced using CHARMM-GUI (Brooks et al., 2009; Jo et al., 2008; Lee et al., 2016) by phosphorylating specific threonine residues resulting in two variants pT549 (phosphorylation at Thr549) and pT556/pT549 (phosphorylation at Thr556 and Thr549 respectively). The protein was then solvated in a rectangular box with a padding size of 10Å utilizing an explicit TIP3P water model for solvation. The system was then neutralized with Potassium (K^+^) and Chloride (Cl^-^) ions and the resultant coordinate and topology files were used as input files for carrying out molecular dynamics (MD) simulations in GROMACS version 2020.7 (Abraham et al., 2015; Bekker et al., 1993; Berendsen et al., 1995; Hess et al., 2008; Lee et al., 2016; Lindahl et al., 2001; Páll et al., 2015; Pronk et al., 2013; Van Der Spoel et al., 2005), compiled with PLUMED 2.9.0 (Bonomi et al., 2019).

The MD simulations were conducted in two phases: an optimization/equilibration phase followed by production runs. The optimization/equilibration process began with energy minimization for 5000 steps using the steepest descent algorithm, applying harmonic restraints of 1000 kJmol^-1^nm^-^² to all protein-heavy atoms to resolve steric clashes or artifacts involving hydrogens. After minimization, the system was heated to 303 K and equilibrated for 125 ps with the same restraints. This minimization and heating cycle were then repeated, but with reduced harmonic restraints of 100 kJmol^-1^nm^-^² to allow for a controlled relaxation of the protein structure. Subsequent equilibration steps consisted of several NVT runs using the Nosé-Hoover thermostat (Evans & Holian, 1985; Hoover, 1985; Nosé, 1984) as temperature control, starting with 100 kJmol^-1^nm^-^² restraints for 125 ps. The restraints were then progressively reduced to 10 kJmol^-1^nm^-^² and to 0.1 kJmol^-1^nm^-^², each for 125 ps, enabling the protein to reach equilibrium. Once optimization/equilibration was complete, production

MD simulations were performed under an NPT ensemble for 50 ns. The temperature was maintained at 303 K using the Nosé-Hoover thermostat, while pressure was regulated by the Berendsen barostat (Berendsen et al., 1984). To ensure statistically reliable results, we ran 10 independent production runs for each system. We used the final, equilibrated structures and velocities from the initial equilibration run as starting points for these production runs.

To assess whether the system had reached equilibration, we calculated the Root Mean Square Deviation (RMSD) of atomic positions over time using VMD 1.9.4 (Humphrey et al., 1996). RMSD was estimated using the backbone atoms of the protein and computed using the equilibrated structure as a reference.

The recently introduced statistical metric Cumulative Variance of atomic Coordinate Fluctuations (CVCF) (Paul et al., 2020) was measured using the following equation:

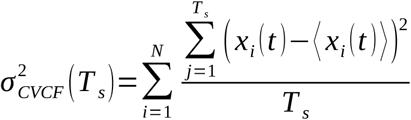

Where *T _s_* refers to the sampling time and *j* refers to all the time steps from the beginning (*j* =1) to the sampling time (*j*=*T_s_*) for which the 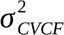 is being calculated. We plot the 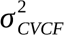 obtained at different *T _s_*. The CVCF was calculated for two subsets of atoms after aligning each subset to an initial structure to eliminate rigid-body translations and rotations. The time step for the CVCF trace was set to 0.1 ns. The two subsets analyzed were the structured protein backbone (Supplementary table 2) and all active site atoms of protein residues surrounding ATP in aPKC pT556+T549 and aPKC pT556. To identify these residues, we modeled ATP-bound aPKC pT556 using AlphaFold-3 (Abramson et al., 2024) and listed all residues within specified cut-off distances of the ATP molecule. The chosen cut-off distances were 3, 5, 7, and 10 Å. The CVCF values for each subset was normalized by the number of atoms to arrive at the per-atom CVCF (total CVCF / respective number of atoms in each subset). Finally, we plotted the individual per-atom CVCF traces for each trajectory and the averaged CVCF trace across all trajectories along with their standard deviations across the full set of trajectories, for both aPKC pT556+T549 and aPKC pT556.

For the structured backbone subset, we analyzed the per-atom CVCF trace by fitting a linear regression to 1-ns intervals of data and calculating the corresponding slopes. To quantify these plateaus better we fitted lines with varying slopes (0.01 – 0.0001 Å²/ns) across the CVCF trajectories computed over 1 ns interval (supp fig. 4C). A slope of zero corresponds to achieving full Boltzmann statistics and the chosen slope thresholds represent variations in atomic positions of less than 0.5-6.0% of typical C–C, C–N, or C–O bond lengths. Varying the slope by two orders of magnitude does not impact our conclusions on statistics such as residence time (Supp fig. 4C).

The RMSD distance of the pT556 and residues of the glycine-rich motif (Supplementary table 1) was calculated from a time when the protein reached a relative thermal equilibrium according to the 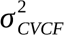 vs *T _s_* plot. The molecular graphics of the protein 3D structures were generated using UCSF ChimeraX ver 1.7.1 (Pettersen et al., 2004).

### Data plotting and statistical analysis

All the quantified data were plotted using R (R Core Team, 2023) and Rstudio (RStudio Team, 2023). The following packages were used for plotting: openxlsx (Schauberger & Alexander, 2024), tidyverse (Wickham et al., 2019), ggplot2 (Wickham, 2016), ggbeeswarm (Clarke et al., 2023), and RColorBrewer (Neuwirth, 2014). The data were compared using statistical tests such as Welch’s t-test, Mann-Whitney U test and Kruskal Wallis test with post hoc using base R and additional packages (asht (Fay, 2023), FSA (Ogle et al., 2023), and dunn.test (Dinno, 2024)) or Past 4.14 software (Hammer et al., 2001). A probability threshold of <0.05 is used determine the significance. The Y-axis of the violin plots is in Log_2_ range for better visualization. The width of the violin at any Y value represents the relative difference in microridge numbers between conditions having that particular length. The box-jitter plots show the median and ranges from 25^th^ and 75^th^ percentile while the error bar represents the 95% confidence of interval. All the numerical data used for plotting the graphs in this study is presented in supplementary data-1. The statistical tests and comparisons conducted on the data sets are shown in supplementary data-2.

## Acknowledgements

We thank Prof. Ullas Kolthur, TIFR, Hyderabad, India for providing HEK293T cell culture line and Prof. Jomon Joseph, NCCS, Pune, India for providing pLgl1/2 (ab59950) antibody used in this article. We also thank Dr. Kalidas Kohale and Boby KV for fish facility and confocal microscope facility maintenance, respectively, and TIFR-DAE (RTI4003: 1303/2/2019/R&D-II/DAE/2079) for funding. R.V. wants to extend his acknowledgement to the Department of Atomic Energy, Government of India for funding support, under Project No. 12-R&D-TFR-5.10-0100.

## Author contribution statement

Performed experiments, S.S., S.M., and S.P.; developed MD Simulation, P.G.; analyzed data, S.S., P.G. S.M., and S.P.; prepared data analysis script, C.S.P.; discovered the phenotype and optimized initial experimental conditions, K.G., G.K.; conceived and conceptualized the project and interpreted the data, S.S., S.M., M.S., and R.V.; secured the funding and supervised the project, M.S. and R.V.; wrote the manuscript, S.S., P.G., R.V., and M.S.

## Declaration of interests

The authors declare no competing interests.

## Figure Legends

**Supplementary Figure 1.**
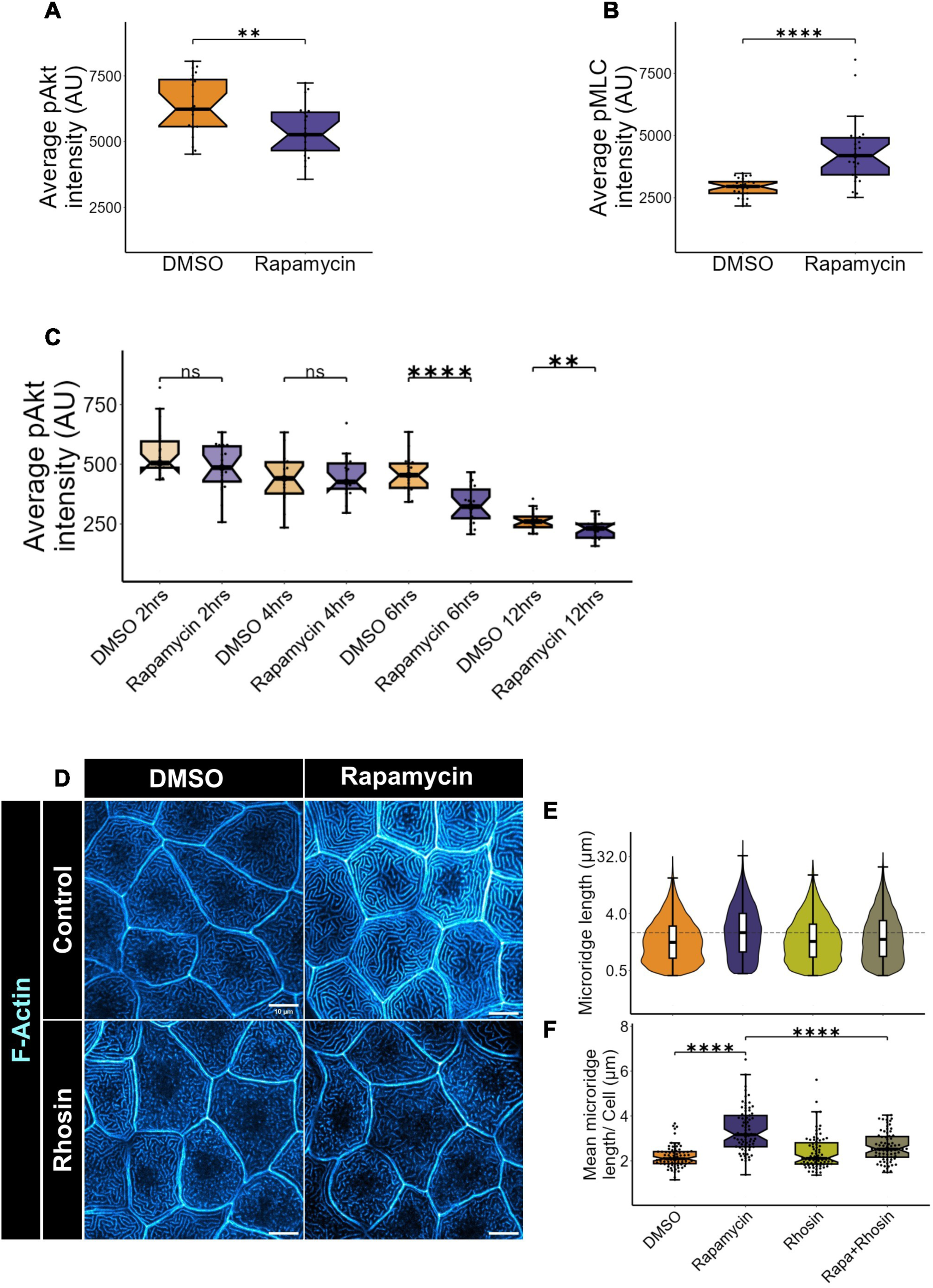
Regulation of microridge formation by mTOR-RhoA axis Box-scatter plot showing the difference in pAkt. (A) and pMLC (B) intensity at 48hpf post 24 hours treatment with rapamycin and DMSO while (C) showing the pAkt level at different time points after rapamycin and DMSO treatment starting at 24 hpf. All three data sets are part of experiments presented in Fig. 1C&D, Fig. 1H&I, and Fig. 3C&D, respectively. Confocal images of peridermal cells stained with rhodamine phalloidin from 48 hpf wildtype embryo (D) treated with vehicle control (DMSO), rapamycin, rhosin, or both starting from 24hpf. The distribution of microridge lengths is plotted in (E) whereas that of mean microridge lengths per cell is plotted in (F). For (A) 20-21 cells from 6 embryos, (B) 25 cells from 5 embryos, and (C) 16-22 cells from 5-6 embryos were analyzed. Experiments (D-F) were performed three times independently and 75 cells from 15 embryos were analyzed. Asterisks denote the significant difference when analyzed using the Mann-Whitney U test for (B) and (F) and student’s t-test for (A) and (C). *ns* (non significant) p>0.05, ** p<0.01, **** p<0.0001. The scale bar equals to 10μm.

**Supplementary Figure 2.**
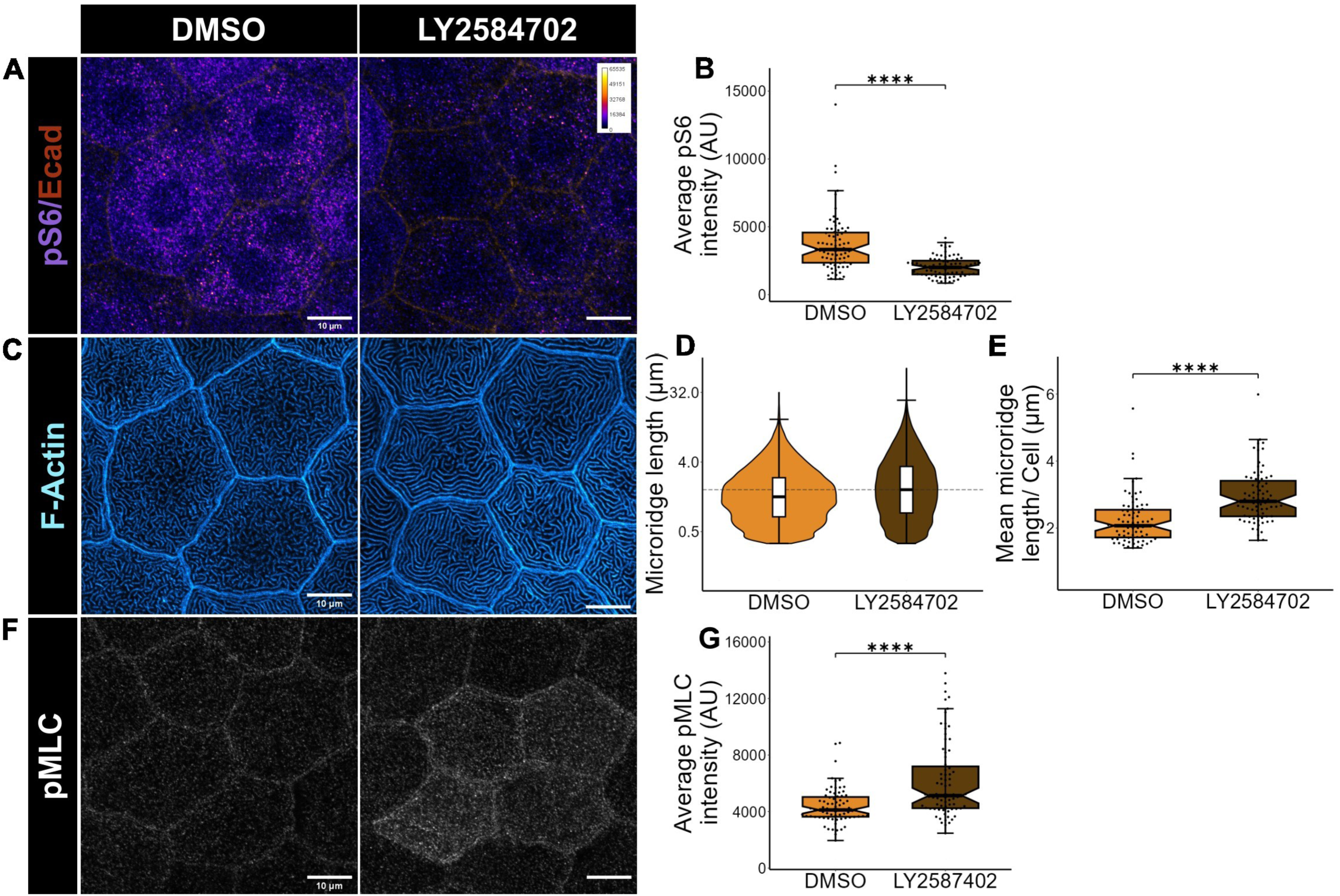
S6K1 inhibition results in elongation of microridges Confocal image of peridermal cells of 48 hpf embryo stained for pS6 (Ser240/244) (A), rhodamine phalloidin (C), or pMLC (Ser19) (F) and treated with LY2584702 or DMSO starting from 24 hpf and quantified for pS6 intensity (B), microridge lengths (D), mean microridge lengths (E), and pMLC intensity (G). The images in (A) are represented in Fire LUT and the calibration bar ranges from 0-65535 indicating lowest to highest intensity. All the experiments were performed three times independently. For all experiments, 75 cells from 15 embryos were analyzed. Asterisks denote the significant difference when analyzed using the Mann-Whitney U test for all the box-jitter plots. **** p<0.0001. The scale bar equals to 10μm.

**Supplementary Figure 3.**
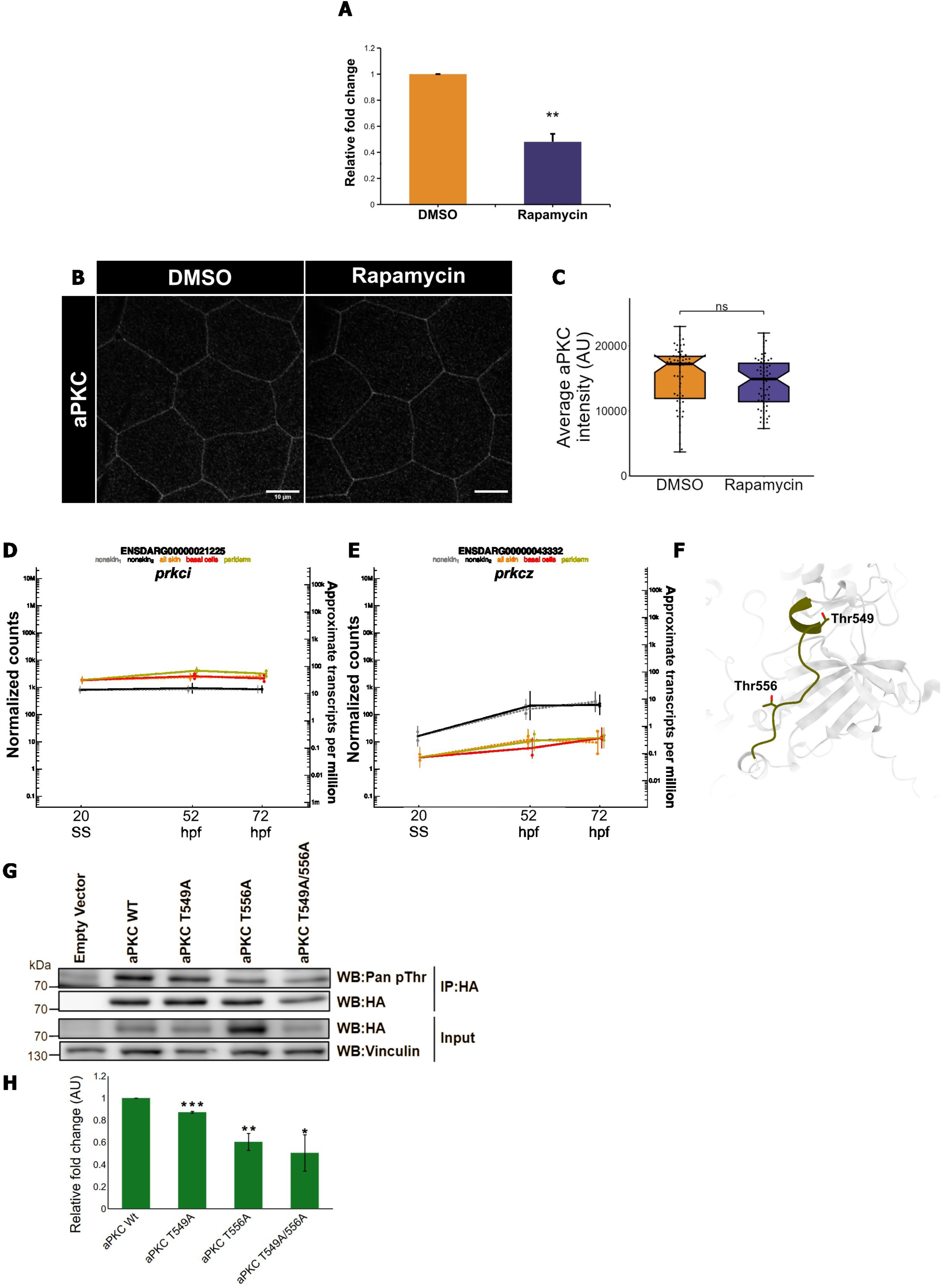
Levels of aPKC protein do not change in the peridermal cells upon mTOR inhibition and aPKC is phosphorylated at thr549 and thr556 sites. (A) qPCR analysis of relative fold change of *prkci* gene (normalized to *eef1a1*) upon rapamycin treatment for 24 hours starting from 24 hpf. (B) aPKC localisation in the peridermal cells of the embryos at 48 hpf treated with rapamycin or DMSO for 24 hours and quantification (C) of aPKC intensity under these two conditions. mRNA expression profiles of *prkci* (D) and *prkcz* (E) in zebrafish embryonic epidermis as per the previous report (Cokus et al., 2019). (F) Alphafold2 model of wildtype aPKC structure highlighting the juxtaposition of Thr549 of TOR Interacting Motif (TIM) and Thr556 of turn motif. Western blot analysis (G) and its quantification (H) for pan phospho Thr level of the mentioned aPKC variants immunoprecipitated using HA antibody. All the experiments were performed three times independently. For data presented in (B) 60-61 cells were analyzed from 18 embryos. Asterisks denote significant differences when analyzed using the student’s t-test in (A, H) or the Mann-Whitney U test for (C). *ns* (non significant) p>0.05, * p<0.05, ** p<0.01, *** p<0.001. The error bars in (H) denotes SEM. The scale bar corresponds to 10μm.

**Supplementary Figure 4.**
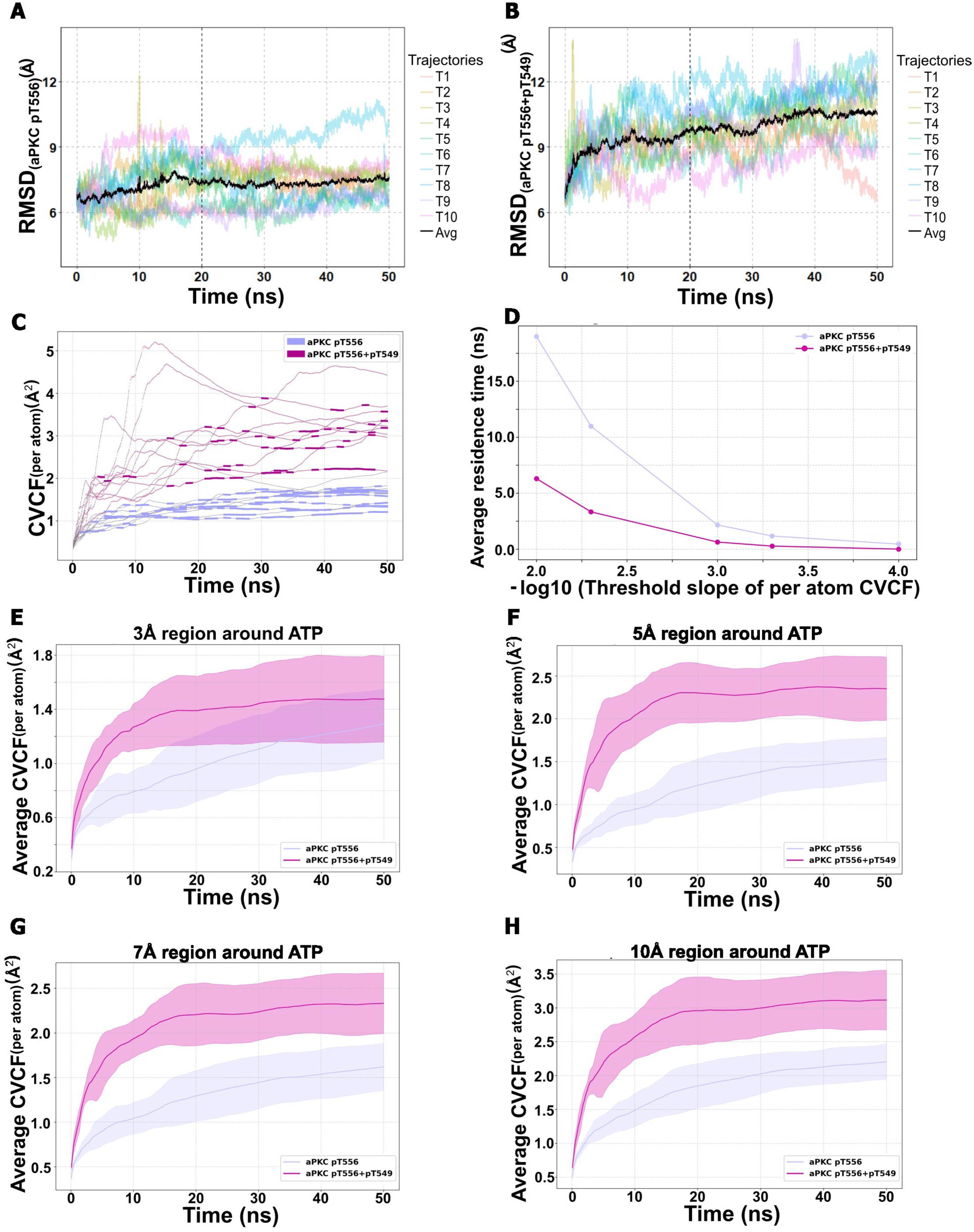
Molecular dynamics simulation analysis shows decreased stability of aPKC pT556+pT549 compared to aPKC pT556 Line plot of RMSD vs Time from MD Simulation of aPKC pT556. (A) and pT556/pT549 (B). The colored lines are from 10 individual iterations and the black line is the average trend line. The line plot of CVCF vs Time (C) of all 10 trajectories from MD Simulation of aPKC pT556 (purple) and pT556/pT549 (magenta) shows plateaus which indicates the metastable stages of the protein. The comparison of these metastable states between aPKC pT556 and aPKC pT556+pT549 is shown in average residence time plot (D). The X-axis has been represented in -Log_10_ for better visualization. The average CVCF trace of the residues surrounding the region of ATP binding cleft of aPKC calculated within the region of 3Å (E), 5Å (F), 7Å (G), and 10Å (H) of ATP.

**Supplementary table 1:**
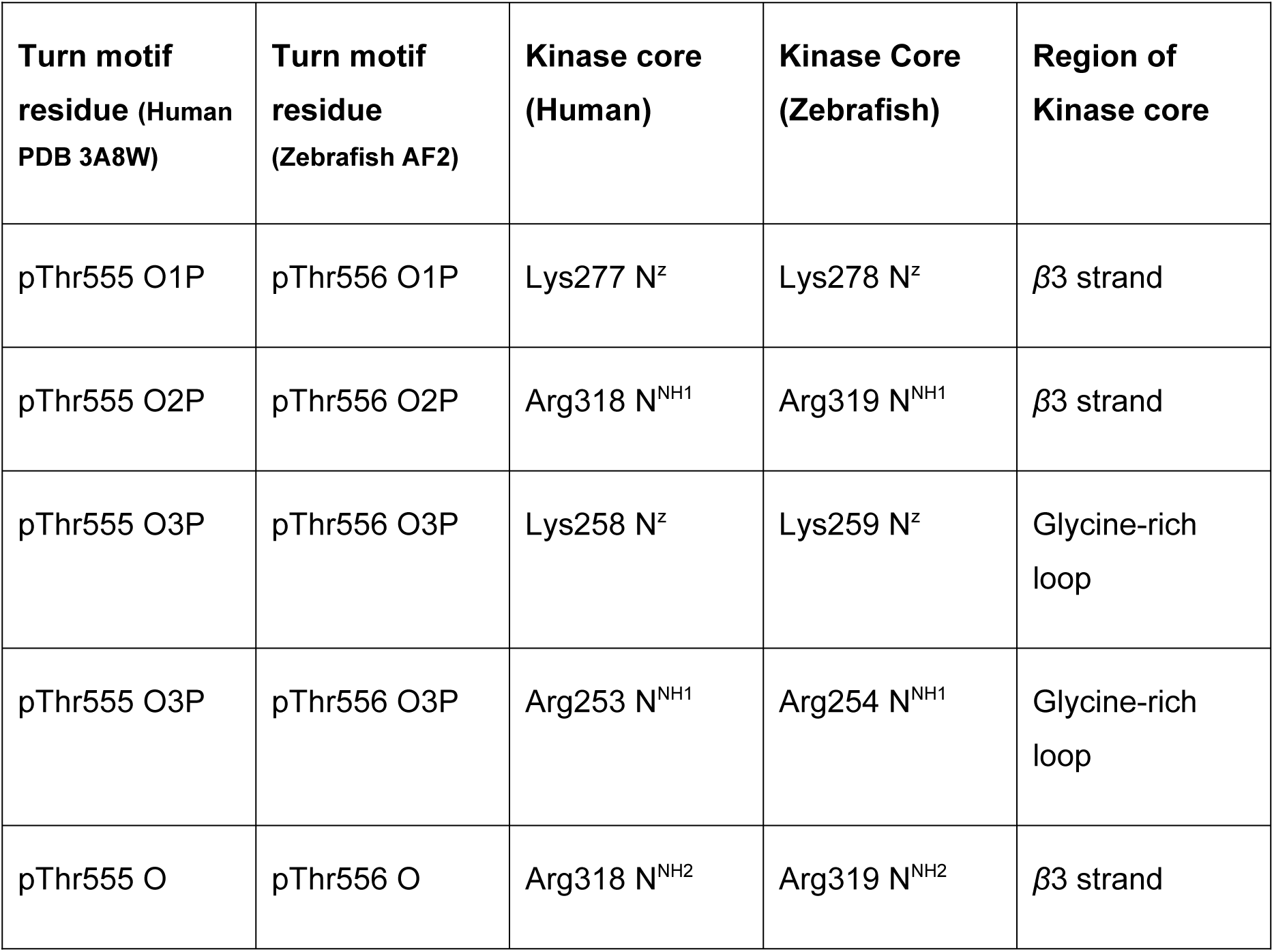
Interaction between the phosphate group of Turn motif (T555/556, human/zebrafish aPKC) and kinase core according to Takimura et al., 2010.

**Supplementary table 2:** A list of structured regions of aPKC wildtype AlphaFold2 structure.

| **Residues** | **Secondary Structures** |
| --- | --- |
| 19 to 25 | Extended Beta Sheet |
| 28 to 34 | Extended Beta Sheet |
| 40 to 51 | Alpha Helix |
| 60 to 64 | Extended Beta Sheet |
| 70 to 73 | Extended Beta Sheet |
| 76 to 89 | Alpha Helix |
| 94 to 99 | Extended Beta Sheet |
| 126 to 131 | Extended Beta Sheet |
| 134 to 139 | Extended Beta Sheet |
| 161 to 164 | Extended Beta Sheet |
| 170 to 172 | Extended Beta Sheet |
| 176 to 178 | 3_10 Helix |
| 243 to 246 | 3_10 Helix |
| 247 to 254 | Extended Beta Sheet |
| 258 to 265 | Extended Beta Sheet |
| 270 to 281 | Extended Beta Sheet |
| 285 to 302 | Alpha Helix |
| 310 to 315 | Extended Beta Sheet |
| 319 to 325 | Extended Beta Sheet |
| 330 to 332 | Extended Beta Sheet |
| 333 to 339 | Alpha Helix |
| 344 to 363 | Alpha Helix |
| 366 to 367 | Extended Beta Sheet |
| 373 to 375 | 3_10 Helix |
| 376 to 378 | Extended Beta Sheet |
| 384 to 386 | Extended Beta Sheet |
| 393 to 394 | Extended Beta Sheet |
| 409 to 411 | 3_10 Helix |
| 414 to 418 | Alpha Helix |
| 424 to 440 | Alpha Helix |
| 454 to 456 | 3_10 Helix |
| 459 to 468 | Alpha Helix |
| 479 to 489 | Alpha Helix |
| 493 to 495 | 3_10 Helix |
| 505 to 510 | Alpha Helix |
| 512 to 514 | 3_10 Helix |
| 519 to 524 | Alpha Helix |
| 546 to 550 | Alpha Helix |
| 560 to 565 | Alpha Helix |
| 568 to 571 | 3_10 Helix |
| 576 to 577 | Extended Beta Sheet |
| 579 to 583 | Alpha Helix |

## Notes

### Competing Interest Statement

The authors have declared no competing interest.

### Summary of Updates

The section titled 'TIM motif phosphorylation leads to decreased interaction between pThr556 of the turn motif and Arg254 of the glycine-rich motif' has been updated with new data. An additional author has been included. Supplemental data added.

