## Supplementary Information for "mTOR signaling governs the formation of epithelial apical projection via S6K1-RhoA and aPKC-Lgl2 axes"

**Supplementary information 1. List of primers used in this study**

| Gene name | Used for | Sequence (5'→ 3') |
|---|---|---|
| aPKC T549E forward | SDM | GATAACTTTGATGCCCAGTTCGAGAACGAG CCCATTCAGCTCACG |
| aPKC T549E reverse | SDM | CGTGAGCTGAATGGGCTCGTTCTCGAACTG GGCATCAAAGTTATC |
| aPKC T549A forward | SDM | CTTTGATGCCCAGTTCGCCAACGAGCCCAT TCA |
| aPKC T549A reverse | SDM | TGAATGGGCTCGTTGGCGAACTGGGCATCA AAG |
| aPKC A122E forward | SDM | ATATACCGGCGGGGAGAGCGACGTTGGAG GAAAC |
| aPKC A122E reverse | SDM | GTTTCCTCCAACGTCGCTCTCCCCGCCGGT ATAT |
| *prkci* forward | qPCR | CAAGGAGTCGAAGGAACG |
| *prkci* reverse | qPCR | CAAGGAGTCGAAGGAACG |
| *eef1a1* forward | qPCR | GAGGCCAGCTCAAACATGG |
| *eef1a1* reverse | qPCR | GCAGAATGGCATCAAGGG |

### Supplementary information 2: Code for microridge analysis

```
/*Clyde S. Pinto
 * 2018
 * ridgeQuant
 *
 *
 * The purpose of this code is to obtain all relavant data with regards to ridge analysis
 * You need to make a closed selection such as a polygon or oval. Square or rectangular selections do work,
but not with directionality.
 * Once the appropriate selection is made save it to the roiManager, usually by pressing "t" or "Ctrl-t" or
"Command-t".
 * Then run the macro and set appropriate parameters and say ok. Then wait till completion.
 */



function arrayBacktoFront(array, index){ //the original array starts at a random place in terms of the actual
ridge ends, this sets the array from an end once the end point is known
	if (index == 0){
		reorderedArray = array;
	}
	else {
		trim = Array.trim(array, (index));
		slice = Array.slice(array,index,lengthOf(array));
		reorderedArray = Array.concat(slice,trim);
	}
return reorderedArray;
}
//Richard Wheeler
function indexOfArray(array, value) { //finds the index in an array
	count=0;
	for (a=0; a<lengthOf(array); a++) {
		if (array[a]==value) {
			count++;
		}
	}
	if (count>0) {
		indices=newArray(count);
		count=0;
		for (a=0; a<lengthOf(array); a++) {
			if (array[a]==value) {
				indices[count]=a;
				count++;
			}
		}
		return indices;
	}
}
//Richard Wheeler
function appendToArray(value, array) { // appends new values to an existing array
	temparray=newArray(lengthOf(array)+1);
	for (i=0; i<lengthOf(array); i++) {
		temparray[i]=array[i];
	}
	temparray[lengthOf(temparray)-1]=value;
	array=temparray;
	return array;
}

function arrayInsert(array, index, value){//adds a new value into the array
	if (index == -1){
		array = Array.concat(value, array);
```

```
		}
		else {
			array2 = Array.trim(array, index+1);
			array3 = Array.slice(array, index+1, lengthOf(array));
			array4 = Array.concat(array2, value);
			array = Array.concat(array4, array3);
		}
		return array;
}

function getLineLength(x3 , y3, x4, y4){//gets the length of the line that is drawn
		return sqrt(pow((x4-x3),2) + pow((y4-y3),2));
}

function deleteFromArray(array, index){//deletes values from an array
		firstPart = Array.trim(array, index);
		h = Array.slice(array, index+1, lengthOf(array));
		array = Array.concat(firstPart,h);
		return array;
}


function variableName("name", counter){//makes a new variable name for each counter of interest
		newName = ("name"+counter);
		return newName;
}

function getMeanDistanceFromCentroid(xArray, yArray, name){

		distanceFromCentroid = 0;
			for (point = 0; point < lengthOf(xArray); point++){
				lineLength = getLineLength(xArray[point], yArray[point], centroidX/width,
centroidY/width);
				distanceFromCentroid += lineLength;
			}
			meanDistanceFromCentroid = width*(distanceFromCentroid/lengthOf(xArray));
			name = meanDistanceFromCentroid;

			return name;
}

function getMaxDistanceFromCentroid(xArray, yArray){

		longestLineLength = 0;
			for (point = 0; point < lengthOf(xArray); point++){
				newLineLength = getLineLength(xArray[point], yArray[point], centroidX/width,
centroidY/width);
				if (newLineLength > longestLineLength){
					longestLineLength = newLineLength;
				}
			}
			return width*longestLineLength;//width multiplication here or (multplying xArray[point] and
yArray[point] while not dividing centroidX and centroidY with width) give the same result
}



//modified from John Lim

function lengthTable(a,b,c,d,e,f,g,h,i,j,k,l,m, n,o,p,q,r,s,t){//generates the length table
		title1="Length";
		title2="["+title1+"]";//This pattern "["+variable+"]" is important for the print command to print in the
appropriate table and columns
```

```
	if (isOpen(title1)){
		print(title2,
a+"\t"+b+"\t"+c+"\t"+d+"\t"+e+"\t"+f+"\t"+g+"\t"+h+"\t"+i+"\t"+j+"\t"+k+"\t"+l+"\t"+m+"\t"+n+"\t"+o+"\t"+p+"\t"+q+
"\t"+r+"\t"+s+"\t"+t);
	}
	else{
		run("Table...", "name="+title1+" width=1000 height=700");
		print(title2, "\\Headings:"+"Cell number"+"\t"+"Line Number"+"\t"+"Length"+"\t"+"Corrected
Length"+"\t"+"Distance of Line from centroid"+"\t"+"Max Distance of Cell Perimeter from
Centroid"+"\t"+"Normalised distance of Line from
centroid"+"\t"+"Tortuosity"+"\t"+"Start"+"\t"+"End"+"\t"+"pixelWidth"+"\t"+"Correction
Factor"+"\t"+"Units"+"\t"+"Genotype"+"\t"+"Treatment"+"\t"+"Dose"+"\t"+"Region"+"\t"+"Time
Point"+"\t"+"Set"+"\t"+"Image Name");
		print(title2,
a+"\t"+b+"\t"+c+"\t"+d+"\t"+e+"\t"+f+"\t"+g+"\t"+h+"\t"+i+"\t"+j+"\t"+k+"\t"+l+"\t"+m+"\t"+n+"\t"+o+"\t"+p+"\t"+q+
"\t"+r+"\t"+s+"\t"+t);
	}
}

function summaryTable(a,b,c,d,e,f,g,h,i,j,k,l,m,o,p,q,r,s,t,u){//generates the summary table
	title1="Summary";
	title2="["+title1+"]";//This pattern "["+variable+"]" is important for the print command to print in the
appropriate table and columns
	if (isOpen(title1)){
		print(title2,
a+"\t"+b+"\t"+c+"\t"+d+"\t"+e+"\t"+f+"\t"+g+"\t"+h+"\t"+i+"\t"+j+"\t"+k+"\t"+l+"\t"+m+"\t"+o+"\t"+p+"\t"+q+"\t"+r+"
\t"+s+"\t"+t+"\t"+u);
	}
	else{
		run("Table...", "name="+title1+" width=1000 height=500");
		print(title2, "\\Headings:"+"Cell"+"\t"+"Area"+"\t"+"Area Fraction(%)"+"\t"+"Ridge
count"+"\t"+"Intersection count"+"\t"+"Ridge count/Area"+"\t"+"Intersection count/Area"+"\t"+"Mean Ridge
Length"+"\t"+"Corrected Mean Ridge Length"+"\t"+"Corrected Mean Ridge Length/Area"+"\t"+"Total Ridge
Length"+"\t"+"Corrected Total Ridge Length"+"\t"+"Corrected Total Ridge
Length/Area"+"\t"+"Genotype"+"\t"+"Treatment"+"\t"+"Dose"+"\t"+"Region"+"\t"+"Time
Point"+"\t"+"Set"+"\t"+"Image Name");
		print(title2,
a+"\t"+b+"\t"+c+"\t"+d+"\t"+e+"\t"+f+"\t"+g+"\t"+h+"\t"+i+"\t"+j+"\t"+k+"\t"+l+"\t"+m+"\t"+o+"\t"+p+"\t"+q+"\t"+r+"
\t"+s+"\t"+t+"\t"+u);
	}
}

function autoLocalThresholdSetting(value, number, parameter_1, parameter_2){//makes a new variable
name for each counter of interest
	run("Auto Local Threshold","method="+value+" radius="+number+" parameter_1="+parameter_1+"
parameter_2="+parameter_2+" white\"");
}

requires("1.49a");
startTime = getTime();



if (isOpen("Results")) {
   selectWindow("Results");
   run("Close");
  }

//colour array is useful for colouring the graph or the ridge lengths
colourArrayDirectionality = newArray("#FCOOOO", "#FF5252", "#FF00D9", "#FC8BEB", "#9B00E3",
"#C240FF", "#0026FF", "#7A8CF0", "#00C3FF", "#B3EBFC", "#00EB66", "#5EFFA4", "#29FF21",
"#9CFF96","#F6FF00", "#FCFF99", "#FF9D00", "#FFCA75", "#FF5500", "#FFB18A");
```

```
colourArrayLength = newArray("#F6FF00", "#FCFF99", "#FF9D00", "#FFCA75", "#FF5500",
"#FFB18A","#FCOOOO", "#FF5252", "#FF00D9", "#FC8BEB", "#9B00E3", "#C240FF", "#0026FF",
"#7A8CF0", "#00C3FF", "#B3EBFC", "#00EB66", "#5EFFA4", "#29FF21", "#9CFF96");


thresholdMethodArray = newArray("Bernsen", "Contrast", "Mean", "Median", "MidGrey",  "Niblack", "Otsu",
"Phansalkar", "Sauvola" );
bifurcationFillColourArray = newArray("red", "green", "blue", "yellow", "orange", "black",
"white","cyan","magenta");
lineFillColourArray = newArray("Auto", "red", "green", "blue", "yellow", "orange", "black",
"white","cyan","magenta");


{  Dialog.create("Settings"); //creates the initial dialogue
	Dialog.addMessage("What condition does this image come from?\neg. Test, Sham, Treated, Drug
Name, WT, Control etc. Leave as NA if unused.");
	Dialog.addString("Genotype:","NA");
	Dialog.addString("Treatment:","NA");
	Dialog.addString("Dose:","NA");
	Dialog.addString("Region:","NA");
	Dialog.addString("Time Point:","NA");
	Dialog.addString("Set: ", "NA");
	Dialog.addMessage("Roi Manager Properties");
			Dialog.addSlider("Ridge Line Width", 0, 5, 2);
			Dialog.addToSameRow();
			Dialog.addChoice("Ridge Line Colour", lineFillColourArray, true);
			Dialog.addSlider("Ridge Bifurcation Width", 0, 5, 2);
	Dialog.addToSameRow();
	Dialog.addChoice("Bifurcation Fill Colour", bifurcationFillColourArray, "black");
	Dialog.addMessage("Auto Local Threshold Settings");
	Dialog.addChoice("Threshold Method", thresholdMethodArray, "Otsu");
	Dialog.addSlider("Threshold Radius", 1, 200, 8);
	Dialog.addNumber("Threshold Parameter 1", 0);
	Dialog.addToSameRow();
	Dialog.addNumber("Threshold Parameter 2", 0);
	Dialog.addCheckbox("Invert", false);
	Dialog.addNumber("Length Correction Factor", 3);
	//Dialog.addMessage("Find feature widths and distance between features?");
	//Dialog.addCheckbox("Find width and distance", false);
	Dialog.addMessage("Run the Directionality Plugin?");
	Dialog.addCheckbox("Run Directionality", false);
	Dialog.addMessage("Get desctiptions of result headings?");
	Dialog.addCheckbox("Get Legend", false);
	Dialog.show();
	genotype = Dialog.getString();
	treatment = Dialog.getString();
	dose = Dialog.getString();
	region = Dialog.getString();
	timepoint = Dialog.getString();
	setNo = Dialog.getString();
	lineWidth = Dialog.getNumber();
	lineFillColour = Dialog.getChoice();
	bifurcationWidth = Dialog.getNumber();

	bifurcationFillColour = Dialog.getChoice();
	thresholdMethod = Dialog.getChoice();
	thresholdRadius = Dialog.getNumber();
	thresholdParam1 = Dialog.getNumber();
	thresholdParam2 = Dialog.getNumber();
	invertAnswer = Dialog.getCheckbox();
	correctionFactor = Dialog.getNumber();
	//widthFinderAnswer = Dialog.getCheckbox();
	directionalityAnswer = Dialog.getCheckbox();
```

```
        legendAnswer = Dialog.getCheckbox();
        }


setBatchMode(true);


numberOfCells = roiManager("Count");
if (numberOfCells==0) {
    exit("Please make a polygon selection and save it in the ROI Manager.");
}
else {
        for (ridgeTest = 0; ridgeTest < numberOfCells; ridgeTest++){
                roiManager("select", ridgeTest);
                if (selectionType>=4 ) {
                        exit("Only Closed Selections Required.");
                }
                else {
                        run("Interpolate", "interval=1 smooth");
                        roiManager("Add");
                }
        }
}




//setBatchMode(true);


if (directionalityAnswer == true){
        List.setCommands;
   if (List.get("Directionality")=="") {
     exit( "Directionality plugin is not installed") ;
   }
   for (ridgeTest = 0; ridgeTest < numberOfCells; ridgeTest++){
                roiManager("select", ridgeTest);
                if (selectionType>=4 || selectionType == 0) {
                        exit("Non-Rectanglular, Non-Square, Closed Selection Required for Directionality.");
                }
        }
}


image = getTitle();//gets the name of the open image
Id = getImageID();//gets the image ID
//run("8-bit");//for files not 8-bit
getVoxelSize(width, height, depth, unit);
run("Set Measurements...", "area centroid center area_fraction display redirect=None decimal=5");//sets the
required measurements

//numberOfCells = roiManager("Count");//here the main macro begins the numberOfCells counter is for the
total number of cells whose roi's have been marked
for (findCellCounter = 0; findCellCounter < numberOfCells; findCellCounter++){//for renaming the roi's as cell
number #
        roiManager("Select", findCellCounter);
        roiManager("Rename", "cell "+(findCellCounter+1)+" roi no. "+(findCellCounter+1));
}
selectImage(Id);
//The next part results in time saving because thresholding and processing is not performed on the whole
image but only on the selected cells
//by making a new image with just the selections present
/*roiManager("Deselect");
```

```
roiManager("Show All");
roiManager("Show None");
roiManager("select", 0);
roiManager("Deselect");
roiManager("Show All");
roiManager("Show None");
run("Duplicate...", " ");*/

//justCellRoiImage = getImageID();
/*justCellRoiImage is an arbitrary name for the new duplicated image
this image will be used to copy the cell selections in the roiManager and clear all the non roi regions
obtaining such an image saves time during the autothresholding step*/

overlappedCellArray = newArray();
//below code identifies if cell selections are overlapping and saves their roiManager index in the
overlappedCellArray for future reference
//if cell selections overlap it creates issues with the code giving an error, by knowning which cells overlap we
can prevent those errors.
for (m=0;m<numberOfCells;m++){//adapted from Jerome http://forum.imagej.net/t/faster-alternative-to-roi-
contains-for-overlapping-rois/2400/2
	for (n=0;n<numberOfCells;n++){
	roiManager('select',newArray(m,n));
	roiManager("AND");
		if ((m!=n)&&(selectionType>-1)) {
			overlappedCellArray = appendToArray(m, overlappedCellArray);
			overlappedCellArray = appendToArray(n, overlappedCellArray);
		}
	}
}
/*the overlapped cell array has each index possibly multiple times, this below code ensures that only one
value of each index occurs
a new array is created named printOverlappedCellArray which includes these index values + 1 to inform the
user at the end which cells were overlapping
so that they know that ridges or parts of ridges in the overlapping parts will be present twice in the results,
once for each cell*/
Array.sort(overlappedCellArray);
for (i = 0; i< lengthOf(overlappedCellArray)-1; i++){
	if (overlappedCellArray[i] == overlappedCellArray[i+1]){
		overlappedCellArray = deleteFromArray(overlappedCellArray, i);
		i--;
	}
}

printOverlappedCellArray = newArray();
for (i = 0; i<lengthOf(overlappedCellArray);i++) {
	printOverlappedCellArray = appendToArray(overlappedCellArray[i]+1, printOverlappedCellArray);
}

startimage = Id;
//Id = justCellRoiImage;

interpolatedNumberOfCells = roiManager("Count");

for (findCellCounter = numberOfCells; findCellCounter < interpolatedNumberOfCells; findCellCounter++){//for
renaming the roi's as cell number #
	roiManager("Select", findCellCounter);
	roiManager("Rename", "interpolated cell "+(findCellCounter-numberOfCells+1)+" roi no. "+
(findCellCounter+1));
	roiManager("Set Color", "blue");
}
```

```
for (cellCounter = numberOfCells; cellCounter < interpolatedNumberOfCells; cellCounter++){//cellCounter is
a counter for cell in roi manager
		showProgress(-(cellCounter-numberOfCells)/(interpolatedNumberOfCells - numberOfCells));
		selectImage(Id);

		roiManager("Select", cellCounter);
		cellSliceNumber = getSliceNumber();


		roiManager("Deselect");
		roiManager("Show All");
		roiManager("Show None");

		selectImage(Id);
		setSlice(cellSliceNumber);

		run("Duplicate...", "title=Duplicate");
		selectWindow("Duplicate");

		roiManager("Select", cellCounter);

		setBackgroundColor(0, 0, 0);
		run("Clear Outside");
		run("8-bit");
		autoLocalThresholdSetting(thresholdMethod, thresholdRadius, thresholdParam1,
thresholdParam2);
		setBackgroundColor(0, 0, 0);
		run("Clear Outside");

		if (invertAnswer == false){
			run("Invert", "slice");
		}
		roiManager("Deselect");
		roiManager("Show All");
		roiManager("Show None");

		run("Duplicate...", "title=Thresholded");
		thresholdedJustCellRoiImageId = getImageID();//this image is essential for obtaining the
area fraction(density) and area measurements
		//The process till the next for loop is adapted from a macro for fingerprint analysis
https://mycarta.wordpress.com/2011/12/14/an-example-of-forensic-image-processing-in-imagej/
		selectWindow("Duplicate");
		run("Smooth");
		run("Duplicate...", "title=Skeleton");
		run("Make Binary");
		run("Skeletonize");
		run("32-bit");
		run("Divide...", "value=255.000");
		run("Enhance Contrast", "saturated=0 normalize");
		run("Duplicate...", "title=Convolution");
		run("Convolve...", "text1=[1 1 1\n1 1 1\n1 1 1\n] stack");
//newline
		if (isOpen("Minutiae")){
			selectWindow("Minutiae");
			run("Close");
		}

		imageCalculator("Multiply create 32-bit", "Skeleton","Convolution");
		selectWindow("Result of Skeleton");
		rename("Minutiae");
		minutiaeImageId = getImageID();

		selectWindow("Skeleton");
```

```
			close();
			selectWindow("Convolution");
			close();
			selectWindow("Duplicate");
			close();
		//}

		cell = (cellCounter-numberOfCells+1); // this variable cell is used for table generation
		selectImage(thresholdedJustCellRoiImageId);
		run("Clear Results");
		roiManager("Select", cellCounter);
		roiManager("Measure");
		print("Analyzing: cell: ", cell, "; Image: ", image);
		areaFraction = getResult("%Area", 0);

		selectImage(minutiaeImageId);
		roiManager("Select", cellCounter);
		area = getResult("Area", 0);
		centroidX = getResult("X", 0);
		centroidY = getResult("Y", 0);

		getSelectionCoordinates(xCellPerimeter, yCellPerimeter);
		maxDistanceOfCellPerimeterFromCentroid =
getMaxDistanceFromCentroid(xCellPerimeter,yCellPerimeter);

		setThreshold(2,3);//This threshold level gives us endpoint+body of the ridge
		priorToPerimeter = roiManager("Count");
		run("Analyze Particles...", "add slice");
		postPerimeter = roiManager("Count"); //postPerimeter - priorToPerimeter = number of ridges in a cell
		selectImage(minutiaeImageId);
		setThreshold(4,6);//This threshold level gives us the bifurcation points of the ridge
		run("Analyze Particles...", "add slice");
		postBifurcation = roiManager("Count"); // postBifurcation - postPerimeter = number of bifurcations in
a cell

		//converting the analyze particle perimeter selection into a line
		selectImage(minutiaeImageId);
		setThreshold(0.5,5);//all values come within this range, but resets the previous thresholding
		wait(500);

		straightLineStartXArray = newArray();
		straightLineEndXArray = newArray();
		straightLineStartYArray = newArray();
		straightLineEndYArray = newArray();
		straightLineRoiArray = newArray();

		meanLength = 0;
		correctedMeanLength = 0;
		totalLength = 0;
		correctedTotalLength = 0;

		for (ridgeParticlePerimeterRoiCounter = priorToPerimeter; ridgeParticlePerimeterRoiCounter <
postPerimeter; ridgeParticlePerimeterRoiCounter++){

			selectImage(minutiaeImageId);
			roiManager("Select", ridgeParticlePerimeterRoiCounter);
			getSelectionCoordinates(ridgeXArray, ridgeYArray);

			for (i=0;i<lengthOf(ridgeXArray);i++){ //i is the 1st ridge coordinate counter
				/*This loop helps to find a start point of the ridge, the ridge arrays need not start at
one end of the ridge, by finding where
				 the end is by looking at neighbouring pixels for intensity of 2 or >=4 we can find the
first end point which we call the start point
```

```
				 if there is no such start point (neighbourPixelIndex = -1)because the ridge is a circle.
				 The variable "neighbourPixelIndex" gets the array index value of the first point that
is a neighbour of a pixel that has a value of 2 or >=4*/
				neighbourPixelIndex = -1;
				if ((getPixel(ridgeXArray[i],ridgeYArray[i])==2)||
(getPixel(ridgeXArray[i],ridgeYArray[i])>=4)){
					neighbourPixelIndex = i;
					i = lengthOf(ridgeXArray);
				}
				else if ((getPixel(ridgeXArray[i]-1,ridgeYArray[i]-1)==2)||
(getPixel(ridgeXArray[i]-1,ridgeYArray[i]-1)>=4)){
					neighbourPixelIndex = i;
					i = lengthOf(ridgeXArray);
				}
				else if ((getPixel(ridgeXArray[i]-1,ridgeYArray[i])==2)||
(getPixel(ridgeXArray[i]-1,ridgeYArray[i])>=4)){
					neighbourPixelIndex = i;
					i = lengthOf(ridgeXArray);
				}
				else if ((getPixel(ridgeXArray[i],ridgeYArray[i]-1)==2)||
(getPixel(ridgeXArray[i],ridgeYArray[i]-1)>=4)){
					neighbourPixelIndex = i;
					i = lengthOf(ridgeXArray);
					}
				else {
					neighbourPixelIndex = -1;
				}
			}

			if (neighbourPixelIndex == -1){
				//using a fix for circlular selections to make jagged lines smooth thus reducing length
and making it more accurate
				for (i=0; i<(lengthOf(ridgeXArray)-1); i++){
					if (ridgeXArray[i] == ridgeXArray[i+1]||ridgeYArray[i] == ridgeYArray[i+1]) {
						ridgeXArray = deleteFromArray(ridgeXArray, i+1);
						ridgeYArray = deleteFromArray(ridgeYArray, i+1);
					}
				}
				ridgeXArray = appendToArray(ridgeXArray[0], ridgeXArray);
				ridgeYArray = appendToArray(ridgeYArray[0], ridgeYArray);
				makeSelection("polyline", ridgeXArray, ridgeYArray);
				run("Interpolate", "interval=1 smooth");// smoothens the line
				roiManager("Add");//adds a circle
				//isCircle = true;//lineStartPointValue = "Circle";
				//End = false;//lineEndPointValue = "Circle";

			}

			else {
				reorderedXArray = arrayBacktoFront(ridgeXArray, neighbourPixelIndex);//reordering
the array to begin at the start point neighbourPixelIndex
				reorderedYArray = arrayBacktoFront(ridgeYArray, neighbourPixelIndex);

				xArrayOfDuplicates = newArray();//array with only the duplicate points
				yArrayOfDuplicates = newArray();
				for (m=0; m < lengthOf(reorderedXArray); m++){//duplicate points in the array are
corner points through which we want to draw the line, if a point comes twice in the
					// reorderedXArray or reorderedYArray arrays it is a duplicate point. The
counter m is the index in the reorderedArray
					for (n = m+1; n < lengthOf(reorderedXArray); n++){  //n = m+1 so that the m
value itself does not recognize itself and only duplicates are detected
						if (reorderedXArray[m]==reorderedXArray[n]){
							if (reorderedYArray[m] == reorderedYArray[n]){
```

```
                                        xArrayOfDuplicates =
appendToArray(reorderedXArray[m], xArrayOfDuplicates);
                                        yArrayOfDuplicates =
appendToArray(reorderedYArray[m], yArrayOfDuplicates);
                                    }
                                }
                            }
                        }
                        endPointX = newArray();//these arrays will hold the coordinates of the ends of the
ridge
                        endPointY = newArray();

                        for (i=0;i<lengthOf(ridgeXArray);i++){//i is the index in ridgeXArray
                            if ((getPixel(ridgeXArray[i],ridgeYArray[i])==2)||
(getPixel(ridgeXArray[i],ridgeYArray[i])>=4)){
                                endPointX = appendToArray(ridgeXArray[i], endPointX);
                                endPointY = appendToArray(ridgeYArray[i], endPointY);
                            }
                            else if ((getPixel(ridgeXArray[i]-1,ridgeYArray[i]-1)==2)||
(getPixel(ridgeXArray[i]-1,ridgeYArray[i]-1)>=4)){
                                endPointX = appendToArray(ridgeXArray[i], endPointX);
                                endPointY = appendToArray(ridgeYArray[i], endPointY);
                            }
                            else if ((getPixel(ridgeXArray[i]-1,ridgeYArray[i])==2)||
(getPixel(ridgeXArray[i]-1,ridgeYArray[i])>=4)){
                                endPointX = appendToArray(ridgeXArray[i], endPointX);
                                endPointY = appendToArray(ridgeYArray[i], endPointY);
                            }
                            else if ((getPixel(ridgeXArray[i],ridgeYArray[i]-1)==2)||
(getPixel(ridgeXArray[i],ridgeYArray[i]-1)>=4)){
                                endPointX = appendToArray(ridgeXArray[i], endPointX);
                                endPointY = appendToArray(ridgeYArray[i], endPointY);
                                }
                            else {
                                neighbourPixelIndex = -1;
                            }
                        }

                        for (m = 0 ; m < lengthOf(endPointX); m++){//m is the index in the endPoint arrays
                            for (n = m + 1; n< lengthOf(endPointX); n++){//n = m+1
                                if (endPointX[m] == endPointX[n]){
                                    if (endPointY[m] == endPointY[n]){
                                        endPointX = deleteFromArray(endPointX, n);
                                        endPointY = deleteFromArray(endPointY, n);
                                    }
                                }
                            }
                        }

                        /*In order to recognise the two parts of the endpoint arrays that signifiy the co-
ordinates at the start of the ridge and those at the end of the ridge
                         * I have chosen to select the index 0 of the endpoint arrays as the 'start' point. If a
co-ordinate in the endpoint arrays is more than 1 pixel distance
                         * away from the start point they are considered as part of the 'end' group of co-
ordinates.
                         */

                        startXArray = newArray();//array having the start points
                        startYArray = newArray();
                        endXArray = newArray();//array having the end points
                        endYArray = newArray();
                        for (i = 1; i < lengthOf(endPointX); i++){
```

```
                    if ((endPointX[i] == (endPointX[0]+1))||(endPointX[i] == (endPointX[0]-1))||
(endPointX[i] == (endPointX[0]))){
                        if ((endPointY[i] == (endPointY[0]+1))||(endPointY[i] ==
(endPointY[0]-1))||(endPointY[i] == (endPointY[0]))){
                            startXArray = appendToArray(endPointX[i], startXArray);
                            startYArray = appendToArray(endPointY[i], startYArray);
                        }
                        else {
                        endXArray = appendToArray(endPointX[i], endXArray);
                        endYArray = appendToArray(endPointY[i], endYArray);
                        }
                    }
                    else {
                        endXArray = appendToArray(endPointX[i], endXArray);
                        endYArray = appendToArray(endPointY[i], endYArray);
                    }
                }

                startXArray = appendToArray(endPointX[0], startXArray);//Adds the 0 position of the
endPointX array to the startXArray array
                startYArray = appendToArray(endPointY[0], startYArray);//Adds the 0 position of the
endPointY array to the startYArray array

                //testing to make lines
                /*ArrayOfDuplicates contains the duplicate points, but doesn't contain the first and
last point
                in order to find the best start and end points we want to find that point in the
startArrays and endArrays that is
                furthest away from the first and last points respectively in the ArrayOfDuplicates*/

                //In this instance the ArrayOfDuplicates == 0; or is empty so the ridge is a straight
line.
                if (lengthOf(xArrayOfDuplicates) == 0) {//xArrayOfDuplicates is the array that
contains duplicate points for line joining
                    if (lengthOf(endXArray)==0 ){
                        valueA = 0;
                        for (m = 0; m < lengthOf(startXArray); m++){  //m is the index in the
startArrays
                            for (n = m+1; n<lengthOf(startXArray); n++){//n is m+1
                                l =
getLineLength(startXArray[m],startYArray[m],startXArray[n],startYArray[n]);
                                if (valueA == 0){
                                    valueA = l;
                                    bestStart = m;
                                    bestEnd = n;
                                }
                                else if (valueA < l){
                                    valueA = l;
                                    bestStart = m;
                                    bestEnd = n;
                                }
                                else {
                                    valueA = valueA;
                                }
                            }
                        }


    makeLine(startXArray[bestStart],startYArray[bestStart],startXArray[bestEnd],startYArray[bestEnd]);//
this make a line a single pixel
                    roiManager("Add");

                    /*if ( ridgeParticlePerimeterRoiCounter == 41){
```

```
					print ("b");
					/
				}*/
			}

		else if (lengthOf(endXArray) != lengthOf(startXArray)) {
			instartx = 0;
			inendx = 0;
			instarty = 0;
			inendy = 0;

			if (lengthOf(endXArray) < lengthOf(startXArray)) {
				for (r = 0; r< lengthOf(endXArray); r++) {
					for (s = 0; s < lengthOf(startXArray);s++) {
						if (endXArray[r] == startXArray[s]) {
							instartx = startXArray[s];
							inendx = endXArray[r];
							instarty = startYArray[s];
							inendy = endYArray[r];
							straightLineStartXArray =
appendToArray((startXArray[0]+startXArray[1])/2, straightLineStartXArray);
							straightLineEndXArray =
appendToArray((startXArray[0]+startXArray[1])/2, straightLineEndXArray);
							straightLineStartYArray =
appendToArray(instarty, straightLineStartYArray);
							straightLineEndYArray =
appendToArray(inendy, straightLineEndYArray);
						}
						else if (endYArray[r] == startYArray[s]) {
							instartx = startXArray[s];
							inendx = endXArray[r];
							instarty = startYArray[s];
							inendy = endYArray[r];
							straightLineStartXArray =
appendToArray(instartx, straightLineStartXArray);
							straightLineEndXArray =
appendToArray(inendx, straightLineEndXArray);
							straightLineStartYArray =
appendToArray((startYArray[0]+startYArray[1])/2, straightLineStartYArray);
							straightLineEndYArray =
appendToArray((startYArray[0]+startYArray[1])/2, straightLineEndYArray);
						}
					}
				}
			}
			else {
				for (r = 0; r< lengthOf(startXArray); r++) {
					for (s = 0; s < lengthOf(endXArray);s++) {
						if (endXArray[s] == startXArray[r]) {
							instartx = startXArray[r];
							inendx = endXArray[s];
							instarty = startYArray[r];
							inendy = endYArray[s];
							straightLineStartXArray =
appendToArray((endXArray[0]+endXArray[1])/2, straightLineStartXArray);
							straightLineEndXArray =
appendToArray((endXArray[0]+endXArray[1])/2, straightLineEndXArray);
							straightLineStartYArray =
appendToArray(instarty, straightLineStartYArray);
							straightLineEndYArray =
appendToArray(inendy, straightLineEndYArray);

						}
```

```
							else if (endYArray[s] == startYArray[r]) {
								instartx = startXArray[r];
								inendx = endXArray[s];
								instarty = startYArray[r];
								inendy = endYArray[s];
								straightLineStartXArray =
appendToArray(instartx, straightLineStartXArray);
								straightLineEndXArray =
appendToArray(inendx, straightLineEndXArray);
								straightLineStartYArray =
appendToArray((endYArray[0]+endYArray[1])/2, straightLineStartYArray);
								straightLineEndYArray =
appendToArray((endYArray[0]+endYArray[1])/2, straightLineEndYArray);
							}
						}
					}
				}

				makeLine(instartx,instarty,inendx,inendy);
				roiManager("Add");
				straightLineRoiArray = appendToArray(roiManager("Count")-1,
straightLineRoiArray);
				/*if ( ridgeParticlePerimeterRoiCounter == 41){
					Array.show("41ARray", startXArray, startYArray,endXArray,
endYArray, straightLineStartXArray,straightLineEndXArray, straightLineStartYArray,straightLineEndYArray);
							print ("a");
							/
						}*/
			}

				else { // for straight line and finding the middle path through it
					/*find the average of the two start coordinates and the two end
coordinates and draw a line using these
					averages as coordinates*/
					meanStartX = 0;
					meanStartY = 0;
					meanEndX = 0;
					meanEndY = 0;
					sumStartX = 0;
					sumStartY = 0;
					sumEndX = 0;
					sumEndY = 0;

					for (o = 0; o < lengthOf(startXArray); o++){
						sumStartX += startXArray[o];
						sumStartY += startYArray[o];
						sumEndX += endXArray[o];
						sumEndY += endYArray[o];
					}
					meanStartX = sumStartX/lengthOf(startXArray);
					meanStartY = sumStartY/lengthOf(startXArray);
					meanEndX = sumEndX/lengthOf(startXArray);
					meanEndY = sumEndY/lengthOf(startXArray);

					straightLineStartXArray = appendToArray(meanStartX,
straightLineStartXArray);
					straightLineEndXArray = appendToArray(meanEndX,
straightLineEndXArray);
					straightLineStartYArray = appendToArray(meanStartY,
straightLineStartYArray);
					straightLineEndYArray = appendToArray(meanEndY,
straightLineEndYArray);
```

```
                                        if (startXArray[0] == endXArray[0] || startYArray[0] == endYArray[0]||
lengthOf(endXArray)==1){

        makeLine(startXArray[0],startYArray[0],endXArray[0],endYArray[0]);
                                        }
                                        else {

        makeLine(startXArray[0],startYArray[0],endXArray[1],endYArray[1]);
                                        }
                                        roiManager("Add");
                                        straightLineRoiArray = appendToArray(roiManager("Count")-1,
straightLineRoiArray);
                                }
                        }

                //get the start and end points

                        else {

                                if (lengthOf(endXArray)==0 ){
                                        if (lengthOf(xArrayOfDuplicates)>= 8){//place of change
                                                newXArray = newArray();
                                                newYArray = newArray();

                                                for (m = 0; m < lengthOf(reorderedXArray); m++){
                                                        for (n = 0; n < lengthOf(startXArray);n++){

                                                                if (reorderedXArray[m]==startXArray[n]){
                                                                        if (reorderedYArray[m] ==
startYArray[n]){
                                                                                newXArray =
appendToArray(m, newXArray);
                                                                                newYArray =
appendToArray(m, newYArray);
                                                                        }
                                                                }
                                                        }
                                                }

                                        //now to find the consecutive numbers starting with index 1
                                                startXArray = newArray();
                                                startXArray =
appendToArray(reorderedXArray[newXArray[0]],startXArray);
                                                startYArray = newArray();
                                                startYArray =
appendToArray(reorderedYArray[newYArray[0]],startYArray);
                                                endXArray = newArray();
                                                endYArray = newArray();
                                                counter = 0;

                                                for (i = 1; i < lengthOf(newXArray); i++){
                                                        if (counter == 0){
                                                                if (newXArray[i] == (newXArray[i-1]+1)){
                                                                        startXArray =
appendToArray(reorderedXArray[newXArray[i]], startXArray);
                                                                        startYArray =
appendToArray(reorderedYArray[newYArray[i]], startYArray);
                                                                        counter = 0;
                                                                }
                                                                else {
                                                                        endXArray =
appendToArray(reorderedXArray[newXArray[i]], endXArray);
```

```
                                                endYArray =
appendToArray(reorderedYArray[newYArray[i]], endYArray);
                                                counter = 1;
                                                //print (counter);
                                        }
                                }
                                else if (counter == 1){
                                        if (newXArray[i] == (newXArray[i-1]+1)){
                                                endXArray =
appendToArray(reorderedXArray[newXArray[i]], endXArray);
                                                endYArray =
appendToArray(reorderedYArray[newYArray[i]], endYArray);
                                                counter = 1;
                                        }
                                        else {
                                                startXArray =
appendToArray(reorderedXArray[newXArray[i]], startXArray);
                                                startYArray =
appendToArray(reorderedYArray[newYArray[i]], startYArray);
                                                counter = 0;
                                                //print(newXArray[i]);
                                        }
                                }
                        }

                        valueB = 0;

                        for (i = 0; i < lengthOf(startXArray); i++){
                                testLineLength =
getLineLength(startXArray[i],startYArray[i],xArrayOfDuplicates[0],yArrayOfDuplicates[0]);
                                if (valueB == 0){
                                        valueB = testLineLength;
                                        startIndex = i;
                                }
                                else if (valueB < testLineLength){
                                        valueB = testLineLength;
                                        startIndex = i;
                                }
                                else {
                                        valueB = valueB;
                                }
                        }
                        startPtX = startXArray[startIndex];//gives the start point X
coordinate
                        startPtY = startYArray[startIndex];//gives the start point Y
coordinate

                        valueB = 0;
                        z = lengthOf(xArrayOfDuplicates)-1;
                        for (i = 0; i < lengthOf(endXArray); i++){
                                testLineLength =
getLineLength(endXArray[i],endYArray[i],xArrayOfDuplicates[z],yArrayOfDuplicates[z]);
                                if (valueB == 0){
                                        valueB = testLineLength;
                                        endIndex = i;
                                }
                                else if (valueB < testLineLength){
                                        valueB = testLineLength;
                                        endIndex = i;
                                }
                                else {
                                        valueB = valueB;
                                }
```

```
                                        }
                                        endPtX = endXArray[endIndex];//gives the end point X
coordinate
                                        endPtY = endYArray[endIndex];//gives the end point Y
coordinate


                                        xArrayOfDuplicates = arrayInsert(xArrayOfDuplicates, -1,
startPtX);
                                        yArrayOfDuplicates = arrayInsert(yArrayOfDuplicates, -1,
startPtY);


                                        xArrayOfDuplicates = arrayInsert(xArrayOfDuplicates,
lengthOf(xArrayOfDuplicates), endPtX);
                                        yArrayOfDuplicates = arrayInsert(yArrayOfDuplicates,
lengthOf(yArrayOfDuplicates), endPtY);

                                        for (i=0; i<(lengthOf(xArrayOfDuplicates)-1); i++){
                                                if (xArrayOfDuplicates[i] == xArrayOfDuplicates[i+1])
{

                                                        if ((i-1)!= -1 ){
                                                                r = xArrayOfDuplicates[i]-
xArrayOfDuplicates[i-1];

                                                                if (r > 0) {
                                                                        if
(abs(yArrayOfDuplicates[i]-yArrayOfDuplicates[i+1])>4){
                                                                                xArrayOfDuplicates
= arrayInsert(xArrayOfDuplicates, i, (xArrayOfDuplicates[i]+1));
                                                                                yArrayOfDuplicates
= arrayInsert(yArrayOfDuplicates, i, ((yArrayOfDuplicates[i]+yArrayOfDuplicates[i+1])/2));
                                                                        }
                                                                        else if
(abs(yArrayOfDuplicates[i]-yArrayOfDuplicates[i+1])<3){
                                                                                xArrayOfDuplicates
= arrayInsert(xArrayOfDuplicates, i, (xArrayOfDuplicates[i]+0.33));
                                                                                yArrayOfDuplicates
= arrayInsert(yArrayOfDuplicates, i, ((yArrayOfDuplicates[i]+yArrayOfDuplicates[i+1])/2));
                                                                        }
                                                                        else {
                                                                                xArrayOfDuplicates
= arrayInsert(xArrayOfDuplicates, i, (xArrayOfDuplicates[i]+0.66));
                                                                                yArrayOfDuplicates
= arrayInsert(yArrayOfDuplicates, i, ((yArrayOfDuplicates[i]+yArrayOfDuplicates[i+1])/2));
                                                                        }
                                                                }
                                                                else if (r < 0){
                                                                        if
(abs(yArrayOfDuplicates[i]-yArrayOfDuplicates[i+1])>4){
                                                                                xArrayOfDuplicates
= arrayInsert(xArrayOfDuplicates, i, (xArrayOfDuplicates[i]-1));
                                                                                yArrayOfDuplicates
= arrayInsert(yArrayOfDuplicates, i, ((yArrayOfDuplicates[i]+yArrayOfDuplicates[i+1])/2));
                                                                        }
                                                                        else if
(abs(yArrayOfDuplicates[i]-yArrayOfDuplicates[i+1])<3){
                                                                                xArrayOfDuplicates
= arrayInsert(xArrayOfDuplicates, i, (xArrayOfDuplicates[i]-0.33));
                                                                                yArrayOfDuplicates
= arrayInsert(yArrayOfDuplicates, i, ((yArrayOfDuplicates[i]+yArrayOfDuplicates[i+1])/2));
                                                                        }
```

```
                                                            else {
                                                                    xArrayOfDuplicates
= arrayInsert(xArrayOfDuplicates, i, (xArrayOfDuplicates[i]-0.66));
                                                                    yArrayOfDuplicates
= arrayInsert(yArrayOfDuplicates, i, ((yArrayOfDuplicates[i]+yArrayOfDuplicates[i+1])/2));
                                                            }
                                                    }
                                            }
                                            else if (i-1 < 0){
                                                    xArrayOfDuplicates =
xArrayOfDuplicates;
                                            }
                                    }
                                    else if (yArrayOfDuplicates[i] ==
yArrayOfDuplicates[i+1]) {

                                            if ((i-1)!= -1 ){
                                                    r = yArrayOfDuplicates[i]-
yArrayOfDuplicates[i-1];

                                                    if (r > 0) {
                                                            if
(abs(xArrayOfDuplicates[i]-xArrayOfDuplicates[i+1])>4){
                                                                    yArrayOfDuplicates
= arrayInsert(yArrayOfDuplicates, i, (yArrayOfDuplicates[i]+1));
                                                                    xArrayOfDuplicates
= arrayInsert(xArrayOfDuplicates, i, ((xArrayOfDuplicates[i]+xArrayOfDuplicates[i+1])/2));
                                                            }
                                                            else if
(abs(xArrayOfDuplicates[i]-xArrayOfDuplicates[i+1])<3){
                                                                    yArrayOfDuplicates
= arrayInsert(yArrayOfDuplicates, i, (yArrayOfDuplicates[i]+0.33));
                                                                    xArrayOfDuplicates
= arrayInsert(xArrayOfDuplicates, i, ((xArrayOfDuplicates[i]+xArrayOfDuplicates[i+1])/2));
                                                            }
                                                            else {
                                                                    yArrayOfDuplicates
= arrayInsert(yArrayOfDuplicates, i, (yArrayOfDuplicates[i]+0.66));
                                                                    xArrayOfDuplicates
= arrayInsert(xArrayOfDuplicates, i, ((xArrayOfDuplicates[i]+xArrayOfDuplicates[i+1])/2));
                                                            }
                                                    }
                                                    else if (r < 0){
                                                            if
(abs(xArrayOfDuplicates[i]-xArrayOfDuplicates[i+1])>4){
                                                                    yArrayOfDuplicates
= arrayInsert(yArrayOfDuplicates, i, (yArrayOfDuplicates[i]-1));
                                                                    xArrayOfDuplicates
= arrayInsert(xArrayOfDuplicates, i, ((xArrayOfDuplicates[i]+xArrayOfDuplicates[i+1])/2));
                                                            }
                                                            else if
(abs(xArrayOfDuplicates[i]-xArrayOfDuplicates[i+1])<3){
                                                                    yArrayOfDuplicates
= arrayInsert(yArrayOfDuplicates, i, (yArrayOfDuplicates[i]-0.33));
                                                                    xArrayOfDuplicates
= arrayInsert(xArrayOfDuplicates, i, ((xArrayOfDuplicates[i]+xArrayOfDuplicates[i+1])/2));
                                                            }
                                                            else {
                                                                    yArrayOfDuplicates
= arrayInsert(yArrayOfDuplicates, i, (yArrayOfDuplicates[i]-0.66));
                                                                    xArrayOfDuplicates
= arrayInsert(xArrayOfDuplicates, i, ((xArrayOfDuplicates[i]+xArrayOfDuplicates[i+1])/2));
                                                    }
```

```
                                        }
                                        else {
                                                yArrayOfDuplicates = 
yArrayOfDuplicates;
                                        }
                                }
                                else if (i-1 < 0){
                                        xArrayOfDuplicates = 
xArrayOfDuplicates;
                                }
                        }
                }

                makeSelection("polyline", xArrayOfDuplicates, 
yArrayOfDuplicates);
                /*if ( ridgeParticlePerimeterRoiCounter == 41){
                        print ("e");
                        /
                }*/
                roiManager("Add");
        }

        //end place of change indexfinder

        else {
                valueA = 0;
                for (m = 0; m < lengthOf(startXArray); m++){
                        for (n = m+1; n<lengthOf(startXArray); n++){
                                testLineLength = 
getLineLength(startXArray[m],startYArray[m],startXArray[n],startYArray[n]);
                                if (valueA == 0){
                                        valueA = testLineLength;
                                        bestStart = m;
                                        bestEnd = n;

                                }
                                else if (valueA < testLineLength){
                                        valueA = testLineLength;
                                        bestStart = m;
                                        bestEnd = n;
                                }
                                else {
                                        valueA = valueA;
                                }
                        }
                }
                if (lengthOf(startXArray) <= 1){
                                waitForUser("this ridge will not be analyzed 
properly");
                                //ridgeParticlePerimeterRoiCounter -= 1;
                                makeLine(reorderedXArray[0], 
reorderedYArray[0], reorderedXArray[round(lengthOf(reorderedXArray)/2)], 
reorderedYArray[round(lengthOf(reorderedXArray)/2)]);
                                roiManager("Add");
                                //user correction
                }

                else {
                        //Array.show("", startXArray, startYArray , endXArray, 
endYArray);

    makeLine(startXArray[bestStart],startYArray[bestStart],startXArray[bestEnd],startYArray[bestEnd]);
                roiManager("Add");
```

```
				}

				/*if (ridgeParticlePerimeterRoiCounter == 41){
					print ("c");

				}*/

			}

		}

		else {
			valueB = 0;

			for (i = 0; i < lengthOf(startXArray); i++){
				testLineLength =
getLineLength(startXArray[i],startYArray[i],xArrayOfDuplicates[0],yArrayOfDuplicates[0]);
				if (valueB == 0){
					valueB = testLineLength;
					startIndex = i;
				}
				else if (valueB < testLineLength){
					valueB = testLineLength;
					startIndex = i;
				}
				else {
					valueB = valueB;
				}
			}
			startPtX = startXArray[startIndex];//gives the start point X coordinate
			startPtY = startYArray[startIndex];//gives the start point Y coordinate

			valueB = 0;
			z = lengthOf(xArrayOfDuplicates)-1;
			for (i = 0; i < lengthOf(endXArray); i++){
				testLineLength =
getLineLength(endXArray[i],endYArray[i],xArrayOfDuplicates[z],yArrayOfDuplicates[z]);
				if (valueB == 0){
					valueB = testLineLength;
					endIndex = i;
				}
				else if (valueB < testLineLength){
					valueB = testLineLength;
					endIndex = i;
				}
				else {
					valueB = valueB;
				}
			}
			endPtX = endXArray[endIndex];//gives the start point X coordinate
			endPtY = endYArray[endIndex];//gives the start point Y coordinate



			xArrayOfDuplicates = arrayInsert(xArrayOfDuplicates, -1, startPtX);
			yArrayOfDuplicates = arrayInsert(yArrayOfDuplicates, -1, startPtY);


			xArrayOfDuplicates = arrayInsert(xArrayOfDuplicates,
lengthOf(xArrayOfDuplicates), endPtX);
			yArrayOfDuplicates = arrayInsert(yArrayOfDuplicates,
lengthOf(yArrayOfDuplicates), endPtY);
```

```
                                        for (duplicatesIndex=0;
duplicatesIndex<(lengthOf(xArrayOfDuplicates)-1); duplicatesIndex++){
                                                if (xArrayOfDuplicates[duplicatesIndex] ==
xArrayOfDuplicates[duplicatesIndex+1]) {

                                                        if ((duplicatesIndex-1)!= -1 ){
                                                                r = xArrayOfDuplicates[duplicatesIndex]-
xArrayOfDuplicates[duplicatesIndex-1];

                                                                if (r > 0) {
                                                                        if
(abs(yArrayOfDuplicates[duplicatesIndex]-yArrayOfDuplicates[duplicatesIndex+1])>4){
                                                                                xArrayOfDuplicates =
arrayInsert(xArrayOfDuplicates, duplicatesIndex, (xArrayOfDuplicates[duplicatesIndex]+1));
                                                                                yArrayOfDuplicates =
arrayInsert(yArrayOfDuplicates, duplicatesIndex, ((yArrayOfDuplicates[duplicatesIndex]
+yArrayOfDuplicates[duplicatesIndex+1])/2));
                                                                        }
                                                                        else if
(abs(yArrayOfDuplicates[duplicatesIndex]-yArrayOfDuplicates[duplicatesIndex+1])<3){
                                                                                xArrayOfDuplicates =
arrayInsert(xArrayOfDuplicates, duplicatesIndex, (xArrayOfDuplicates[duplicatesIndex]+0.33));
                                                                                yArrayOfDuplicates =
arrayInsert(yArrayOfDuplicates, duplicatesIndex, ((yArrayOfDuplicates[duplicatesIndex]
+yArrayOfDuplicates[duplicatesIndex+1])/2));
                                                                        }
                                                                        else {
                                                                                xArrayOfDuplicates =
arrayInsert(xArrayOfDuplicates, duplicatesIndex, (xArrayOfDuplicates[duplicatesIndex]+0.66));
                                                                                yArrayOfDuplicates =
arrayInsert(yArrayOfDuplicates, duplicatesIndex, ((yArrayOfDuplicates[duplicatesIndex]
+yArrayOfDuplicates[duplicatesIndex+1])/2));
                                                                        }
                                                                }
                                                                else if (r < 0){
                                                                        if
(abs(yArrayOfDuplicates[duplicatesIndex]-yArrayOfDuplicates[duplicatesIndex+1])>4){
                                                                                xArrayOfDuplicates =
arrayInsert(xArrayOfDuplicates, duplicatesIndex, (xArrayOfDuplicates[duplicatesIndex]-1));
                                                                                yArrayOfDuplicates =
arrayInsert(yArrayOfDuplicates, duplicatesIndex, ((yArrayOfDuplicates[duplicatesIndex]
+yArrayOfDuplicates[duplicatesIndex+1])/2));
                                                                        }
                                                                        else if
(abs(yArrayOfDuplicates[duplicatesIndex]-yArrayOfDuplicates[duplicatesIndex+1])<3){
                                                                                xArrayOfDuplicates =
arrayInsert(xArrayOfDuplicates, duplicatesIndex, (xArrayOfDuplicates[duplicatesIndex]-0.33));
                                                                                yArrayOfDuplicates =
arrayInsert(yArrayOfDuplicates, duplicatesIndex, ((yArrayOfDuplicates[duplicatesIndex]
+yArrayOfDuplicates[duplicatesIndex+1])/2));
                                                                        }
                                                                        else {
                                                                                xArrayOfDuplicates =
arrayInsert(xArrayOfDuplicates, duplicatesIndex, (xArrayOfDuplicates[duplicatesIndex]-0.66));
                                                                                yArrayOfDuplicates =
arrayInsert(yArrayOfDuplicates, duplicatesIndex, ((yArrayOfDuplicates[duplicatesIndex]
+yArrayOfDuplicates[duplicatesIndex+1])/2));
                                                                        }
                                                                }
                                                        }
                                                        else if (duplicatesIndex-1 < 0){
                                                                xArrayOfDuplicates = xArrayOfDuplicates;
```

```
                        }
                    }
                    else if (yArrayOfDuplicates[duplicatesIndex] ==
yArrayOfDuplicates[duplicatesIndex+1]) {

                        if ((duplicatesIndex-1)!= -1 ){
                            r = yArrayOfDuplicates[duplicatesIndex]-
yArrayOfDuplicates[duplicatesIndex-1];

                            if (r > 0) {
                                if
(abs(xArrayOfDuplicates[duplicatesIndex]-xArrayOfDuplicates[duplicatesIndex+1])>4){
                                    yArrayOfDuplicates =
arrayInsert(yArrayOfDuplicates, duplicatesIndex, (yArrayOfDuplicates[duplicatesIndex]+1));
                                    xArrayOfDuplicates =
arrayInsert(xArrayOfDuplicates, duplicatesIndex, ((xArrayOfDuplicates[duplicatesIndex]
+xArrayOfDuplicates[duplicatesIndex+1])/2));
                                }
                                else if
(abs(xArrayOfDuplicates[duplicatesIndex]-xArrayOfDuplicates[duplicatesIndex+1])<3){
                                    yArrayOfDuplicates =
arrayInsert(yArrayOfDuplicates, duplicatesIndex, (yArrayOfDuplicates[duplicatesIndex]+0.33));
                                    xArrayOfDuplicates =
arrayInsert(xArrayOfDuplicates, duplicatesIndex, ((xArrayOfDuplicates[duplicatesIndex]
+xArrayOfDuplicates[duplicatesIndex+1])/2));
                                }
                                else {
                                    yArrayOfDuplicates =
arrayInsert(yArrayOfDuplicates, duplicatesIndex, (yArrayOfDuplicates[duplicatesIndex]+0.66));
                                    xArrayOfDuplicates =
arrayInsert(xArrayOfDuplicates, duplicatesIndex, ((xArrayOfDuplicates[duplicatesIndex]
+xArrayOfDuplicates[duplicatesIndex+1])/2));
                                }
                            }
                            else if (r < 0){
                                if
(abs(xArrayOfDuplicates[duplicatesIndex]-xArrayOfDuplicates[duplicatesIndex+1])>4){
                                    yArrayOfDuplicates =
arrayInsert(yArrayOfDuplicates, duplicatesIndex, (yArrayOfDuplicates[duplicatesIndex]-1));
                                    xArrayOfDuplicates =
arrayInsert(xArrayOfDuplicates, duplicatesIndex, ((xArrayOfDuplicates[duplicatesIndex]
+xArrayOfDuplicates[duplicatesIndex+1])/2));
                                }
                                else if
(abs(xArrayOfDuplicates[duplicatesIndex]-xArrayOfDuplicates[duplicatesIndex+1])<3){
                                    yArrayOfDuplicates =
arrayInsert(yArrayOfDuplicates, duplicatesIndex, (yArrayOfDuplicates[duplicatesIndex]-0.33));
                                    xArrayOfDuplicates =
arrayInsert(xArrayOfDuplicates, duplicatesIndex, ((xArrayOfDuplicates[duplicatesIndex]
+xArrayOfDuplicates[duplicatesIndex+1])/2));
                                }
                                else {
                                    yArrayOfDuplicates =
arrayInsert(yArrayOfDuplicates, duplicatesIndex, (yArrayOfDuplicates[duplicatesIndex]-0.66));
                                    xArrayOfDuplicates =
arrayInsert(xArrayOfDuplicates, duplicatesIndex, ((xArrayOfDuplicates[duplicatesIndex]
+xArrayOfDuplicates[duplicatesIndex+1])/2));
                            }
                            }
                            else {
                                yArrayOfDuplicates =
yArrayOfDuplicates;
                            }
```

```
                                        }
                                        else if (duplicatesIndex-1 < 0){
                                                xArrayOfDuplicates = xArrayOfDuplicates;
                                        }
                                }
                        }

                        makeSelection("polyline", xArrayOfDuplicates, yArrayOfDuplicates);
                        roiManager("Add");
                        /*if ( ridgeParticlePerimeterRoiCounter == 41){
                                        print ("f");
                                        /
                                }*/

                }
            }
        }
}

postLines = roiManager("count");
perimeters = 1;

for (j = priorToPerimeter; j < postPerimeter; j++){
        roiManager("Select", j);
        roiManager("Rename", "cell "+(cell)+" ridge "+(j+1-priorToPerimeter)+" roi no. "+(j+1));
        roiManager("Set Color", "#EAECEE");
        roiManager("Measure");
        perimeters += 1;
}
//print ("cell "+(i +1)+" ridge = "+perimeters);

bifurcations = 1;
for (k = postPerimeter; k< postBifurcation; k++){
        roiManager("Select", k);
        roiManager("Rename", "cell "+(cell)+" bifurcation "+(k-postPerimeter+1)+" roi no. "+(k+1));

        roiManager("Set Color", bifurcationFillColour);
        roiManager("Set Fill Color", bifurcationFillColour);
        roiManager("Set Line Width", bifurcationWidth);
        bifurcations += 1;
}
//print ("cell "+(cellCounter+1)+" bifurcations = "+bifurcations);

lines = 1;
ridgeNumber = 1;
straightLineArrayIndex = 0;
for (k = postBifurcation; k< postLines; k++){
        run("Clear Results");
        roiManager("Select", k);
        roiManager("Rename", "cell "+(cell)+" line "+(k-postBifurcation+1)+" roi no. "+(k+1));

        getSelectionCoordinates(lineX, lineY);

        selectImage(minutiaeImageId);
        lineStartPointValue = 0;
        lineEndPointValue = 0;

        for (i=0;i<1;i++){
                if ((getPixel(lineX[i],lineY[i])==2)||(getPixel(lineX[i],lineY[i])>=4)){
                        if (getPixel(lineX[i],lineY[i])==2){
                                lineStartPointValue = "Endpoint";
                                i > 0;
                        }
```

```
				else {
					lineStartPointValue = "Bifurcation";
					i > 0;
				}
			}
			else if ((getPixel(lineX[i]-1,lineY[i]-1)==2)||(getPixel(lineX[i]-1,lineY[i]-1)>=4)){
				if (getPixel(lineX[i]-1,lineY[i]-1)==2){
					lineStartPointValue = "Endpoint";
					i > 0;
				}
				else {
					lineStartPointValue = "Bifurcation";
					i > 0;
				}
			}
			else if ((getPixel(lineX[i]-1,lineY[i])==2)||(getPixel(lineX[i]-1,lineY[i])>=4)){
				if (getPixel(lineX[i]-1,lineY[i])==2){
					lineStartPointValue = "Endpoint";
					i > 0;
				}
				else {
					lineStartPointValue = "Bifurcation";
					i > 0;
				}
			}
			else if ((getPixel(lineX[i],lineY[i]-1)==2)||(getPixel(lineX[i],lineY[i]-1)>=4)){
				if (getPixel(lineX[i],lineY[i]-1)==2){
					lineStartPointValue = "Endpoint";
					i > 0;
				}
				else {
					lineStartPointValue = "Bifurcation";
					i > 0;
				}
			}
		}
		for (i = (lengthOf(lineX)-1); i<lengthOf(lineX);i++){
			if ((getPixel(lineX[i],lineY[i])==2)||(getPixel(lineX[i],lineY[i])>=4)){
				if (getPixel(lineX[i],lineY[i])==2){
					lineEndPointValue = "Endpoint";
					i != (lengthOf(lineX)-1);
				}
				else {
					lineEndPointValue = "Bifurcation";
					i != (lengthOf(lineX)-1);
				}
			}
			else if ((getPixel(lineX[i]-1,lineY[i]-1)==2)||(getPixel(lineX[i]-1,lineY[i]-1)>=4)){
				if (getPixel(lineX[i]-1,lineY[i]-1)==2){
					lineEndPointValue = "Endpoint";
					i != (lengthOf(lineX)-1);
				}
				else {
					lineEndPointValue = "Bifurcation";
					i != (lengthOf(lineX)-1);
				}
			}
			else if ((getPixel(lineX[i]-1,lineY[i])==2)||(getPixel(lineX[i]-1,lineY[i])>=4)){
				if (getPixel(lineX[i]-1,lineY[i])==2){
					lineEndPointValue = "Endpoint";
					i != (lengthOf(lineX)-1);
				}
				else {
```

```
                        lineEndPointValue = "Bifurcation";
                        i != (lengthOf(lineX)-1);
                    }
                }
                else if ((getPixel(lineX[i],lineY[i]-1)==2)||(getPixel(lineX[i],lineY[i]-1)>=4)){
                    if (getPixel(lineX[i],lineY[i]-1)==2){
                        lineEndPointValue = "Endpoint";
                        i != (lengthOf(lineX)-1);
                    }
                    else {
                        lineEndPointValue = "Bifurcation";
                        i != (lengthOf(lineX)-1);
                    }
                }
            }
            if (lineStartPointValue == 0){
                lineStartPointValue = "Circle";
                lineEndPointValue = "Circle";
            }

            if (lengthOf(straightLineRoiArray)> straightLineArrayIndex){


if (straightLineRoiArray[straightLineArrayIndex] == k){

                    makeLine(straightLineStartXArray[straightLineArrayIndex],
straightLineStartYArray[straightLineArrayIndex],straightLineEndXArray[straightLineArrayIndex],straightLineE
ndYArray[straightLineArrayIndex]);
                    roiManager("Update");

                    //print(straightLineStartXArray[straightLineArrayIndex],
straightLineStartYArray[straightLineArrayIndex],straightLineEndXArray[straightLineArrayIndex],straightLineE
ndYArray[straightLineArrayIndex]);

                    straightLineArrayIndex++;
                }
            }


            run("Interpolate", "interval=1 smooth");
            roiManager("Measure");
            length = getResult("Length",0);

            euclideanDistance = width*getLineLength(lineX[0] , lineY[0], lineX[lengthOf(lineX)-1],
lineY[lengthOf(lineX)-1]);

            tortuosity = length/euclideanDistance;
            getSelectionCoordinates(lineX2, lineY2);

///
            lineName = 0;
            distanceOfLineFromCentroid = getMeanDistanceFromCentroid(lineX2, lineY2, lineName);

            normalisedDistanceOfLineFromCentroid =
distanceOfLineFromCentroid/maxDistanceOfCellPerimeterFromCentroid;


            roiManager("Set Line Width", lineWidth);//important note: changing the line width changes
the pixel positions seemingly

            if (lineFillColour == "Auto"){
                colourArrayNo = 0;
                for (lineColour = 0; lineColour < 9.5; lineColour += 0.5){
```

```
				if (length < lineColour){
					lineColour > 10;
				}
				else {
					colourArrayNo += 1;
				}
			}
			roiManager("Set Color", colourArrayLength[colourArrayNo]);
		}
		else{
			roiManager("Set Color", lineFillColour);
		}

		//cell = (cellCounter+1);
		totalLength += length;

		correctedLength = length + (width*correctionFactor); // correction factor is the addition of
three pixel widths
		correctedTotalLength += correctedLength;

		lengthTable(cell, ridgeNumber, length, correctedLength,
distanceOfLineFromCentroid,maxDistanceOfCellPerimeterFromCentroid,
normalisedDistanceOfLineFromCentroid,  tortuosity, lineStartPointValue, lineEndPointValue, width,
correctionFactor,  unit, genotype, treatment, dose, region, timepoint, setNo, image );
		lines++;
		ridgeNumber++;
	}
	lines = lines -1;
	ridgebyarea = (perimeters-1)/area;
	bifurcationbyarea = (bifurcations-1)/area;
	meanLength = totalLength/lines;

correctedMeanLength = correctedTotalLength/lines;
correctedMeanLengthbyArea = correctedMeanLength/area;
correctedTotalLengthbyArea = correctedTotalLength/area;

	summaryTable(cell,area,areaFraction,perimeters-1,bifurcations-1, ridgebyarea, bifurcationbyarea,
meanLength, correctedMeanLength,correctedMeanLengthbyArea, totalLength,
correctedTotalLength,correctedTotalLengthbyArea, genotype,treatment, dose, region,timepoint, setNo,
image);

}

//Directionality

if (directionalityAnswer == true) {

	selectImage(startimage);
	angle = newArray();
	for (angl = -90; angl < 90; angl++){
		angle = appendToArray(angl, angle);
	}

	Plot.create("Directions", "Direction Angle relative to the horizontal (Degrees)", "Amount");

	colourCounter = 0;
	cellCounterDirectionality = 0;
	directionCellNo = newArray();

	for (directionIndex = 0; directionIndex < numberOfCells; directionIndex++){
		roiManager("Select", directionIndex);
		run("Duplicate...", " ");
		rename("1");
```

```
			run("Directionality", "method=[Local gradient orientation] nbins=180 histogram=-90
display_table");
			list = getList("window.titles");
			for (directionTable=0; directionTable<list.length; directionTable++){
		if (startsWith(list[directionTable], "Directionality")){
						IJ.renameResults(list[directionTable],"Results");
				}
			}

			Direction = newArray();

			Directionfit = newArray();
			for (directionPlot = 0; directionPlot < nResults; directionPlot++){
				directionAngles = getResult("DUP_1", directionPlot);
				directionFitAngles = getResult("DUP_1-fit", directionPlot);
				Direction = appendToArray(directionAngles, Direction);
				Directionfit = appendToArray(directionFitAngles, Directionfit);
			}
		// Need to figure out if there are more than 20 indiceshow to loop it
		if (cellCounterDirectionality < 19){
			Plot.setLineWidth(1);
			Plot.setColor("#D3D3D3");//light grey
			Plot.add("lines", angle, Direction);
			directionCellNo = appendToArray(("Cell "+directionIndex), directionCellNo);
			Plot.setLineWidth(2);
			Plot.setColor(colourArrayDirectionality[colourCounter+1]);
			Plot.add("lines", angle, Directionfit);
			directionCellNo = appendToArray(("Cell "+directionIndex+" fit"), directionCellNo);
			colourCounter += 1;
			cellCounterDirectionality +=1;
			}

		else {
			Plot.setLineWidth(1);
			Plot.setColor("#D3D3D3");//light grey
			Plot.add("lines", angle, Direction);
			directionCellNo = appendToArray(("Cell "+directionIndex), directionCellNo);
			Plot.setLineWidth(2);
			Plot.setColor("red");
			Plot.add("lines", angle, Directionfit);
			directionCellNo = appendToArray(("Cell "+directionIndex+" fit"), directionCellNo);
			colourCounter += 1;
			cellCounterDirectionality +=1;
		}

		//selectWindow("Results");
		//run("Close");
		selectImage(startimage);

		}

		print("\\Clear");

		print ("Cell "+1);
		legendCellNo = 1;
		for (directionLegendIndex = 1; directionLegendIndex <cellCounterDirectionality*2;
directionLegendIndex++){
			logText = getInfo("log");
			if (directionLegendIndex % 2 == 0){
				print ("\\Update:"+logText+ "Cell "+ legendCellNo );
			}
			else {
				print ("\\Update:"+logText+ "Cell "+ legendCellNo + "-fit");
```

```
                    legendCellNo += 1;
                }
            }
            logText = getInfo("log");
            //print (logText);
            Plot.setLegend(logText);
}
//Directionality end....xxxxxxxxx


endTime = getTime();
print("\\Clear");
/*if (isOpen("Results")) {
    selectWindow("Results");
    run("Close");
   } */
print("Processing Time = "+(endTime - startTime)/1000+"s");

if (lengthOf(printOverlappedCellArray) != 0){
            print("Cell selections overlap. The cells that overlap are as follows");
            for (i = 0; i<lengthOf(printOverlappedCellArray);i++){
                        print ("Cell "+ printOverlappedCellArray[i]);
            }
            print ("Please note that ridges or parts of ridges \nin the overlapping regions will be counted twice,
\nonce for each cell selection.");
}
selectImage(startimage);

setBatchMode(false);
selectWindow("Length");
selectImage(startimage);

selectWindow("ROI Manager");
roiManager("Show All without labels");
```
